# The Lysis Cassette of Jumbophage phiKZ

**DOI:** 10.1101/2025.11.11.687635

**Authors:** Prasanth Manohar, Joshua Wan, Gabriella Ganser, Kiersten Sharp, Ry Young

## Abstract

*Pseudomonas* jumbophage phiKZ is prominent because of the proteinaceous nuclear structure that is formed during the infection cycle, conferring resistance of the replicating phage DNA to anti-phage resistance systems. Progress has been made in deciphering the molecular basis of nucleus formation and other unique aspects of phiKZ biology, but the understanding of how it causes host lysis remains minimal. Here we present bioinformatic, physiological and molecular evidence for a “lysis cassette”, the expression of which is necessary and sufficient for the temporally regulated disruption of the host envelope. This cluster contains genes for the usual components of an MGL (multigene lysis) system, including the holin, endolysin, i-spanin and o-spanin. In addition, a fifth gene in the cluster, encoding a cytoplasmic protein, was found to accelerate the timing of holin-mediated lethality when expressed in trans. Evidence is provided that suggests this lysis regulator protein interacts with the cytoplasmic domain of the phiKZ class III holin. In support of this notion, alpha-fold analysis generated a high-confidence structure of a conserved holin-lysis regulator heterodimer complex. Infections at high multiplicity resulted in slower bulk culture lysis profiles than at low multiplicity, suggesting that phiKZ might have a lysis-inhibition system.

## Introduction

*Pseudomonas* phage phiKZ was one of the first jumbo phages (genome >200kbp) discovered (V. N. Krylov & Zhazykov, 1978; Mesyanzhinov et al., 2002). It has been intensively studied because of a number of unique features, especially its proteinaceous “nucleus” that is formed during infection and serves to protect phage replicating DNA from host defense mechanisms (Mendoza et al., 2020). Despite this celebrity, our understanding of its lysis pathway is limited. The tailed (*Caudoviricetes*) phages of Gram-negative hosts generally use a multi-gene lysis (MGL) strategy, with a minimum of 4 proteins: holin, endolysin, i-spanin and o-spanin. The holin is the key to this system, controlling the timing of lysis and thus the fecundity of the infection. Holins are extremely diverse but can be sorted into at least three topology classes, I, II, and III, based on the number and orientation of their transmembrane domains (TMDs) (Fig.1A). After the turn-on of the morphogenesis genes in the infection cycle, the holin accumulates harmlessly in the cytoplasmic membrane (IM) until suddenly permeabilizing it to initiate the lytic process, allowing the endolysin to attack the peptidoglycan (PG) and terminating macromolecular synthesis (Young, 2014). The saltatory transition from harmless accumulation to lethal permeabilization has been designated “triggering” and is thought to reflect the attainment of a critical concentration of holin in the IM. Notably, premature triggering and thus early lysis can be obtained by as little as a 30% reduction in the proton motive force (PMF) by a variety of means (Cahill et al., 2024; Gründling et al., 2001), so the infected cell is poised to undergo lysis even early in the morphogenesis period. In parallel with the holin, the i-spanin and o-spanin accumulate in the IM and OM, respectively, forming heterotetrameric complexes that span the periplasm and are trapped in the meshwork of the PG. After triggering, degradation of the PG by the endolysin liberates the spanin complexes, which are thought to undergo tertiary and quaternary conformational changes to facilitate destruction of the OM by fusing it with the IM (J. Berry et al., 2010a; J. D. Berry et al., 2013; Cahill, Rajaure, O’Leary, et al., 2017). Early genomic analysis of phiKZ revealed the gene for the endolysin, gp*144*, identified as a lytic transglycosylase with a C-terminal cell wall-binding domain, features that were confirmed by biochemical studies with the purified enzyme (Fokine et al., 2008; Miroshnikov et al., 2006) (Fig.1B). However, no gene with sequence similarity to a known holin gene was identified. This led to speculation that phiKZ and similar jumbophages used an uncharacterized strategy for host lysis (V. Krylov et al., 2021). Here we document the existence of a traditional “lysis cassette” including all four MGL genes and verify the annotations by complementation tests and physiological experiments. In addition, we provide evidence that a fifth gene, *143*, in the lysis cassette has functional characteristics that suggest it is a lysis regulator acting through interactions with the holin.

**Figure 1.**
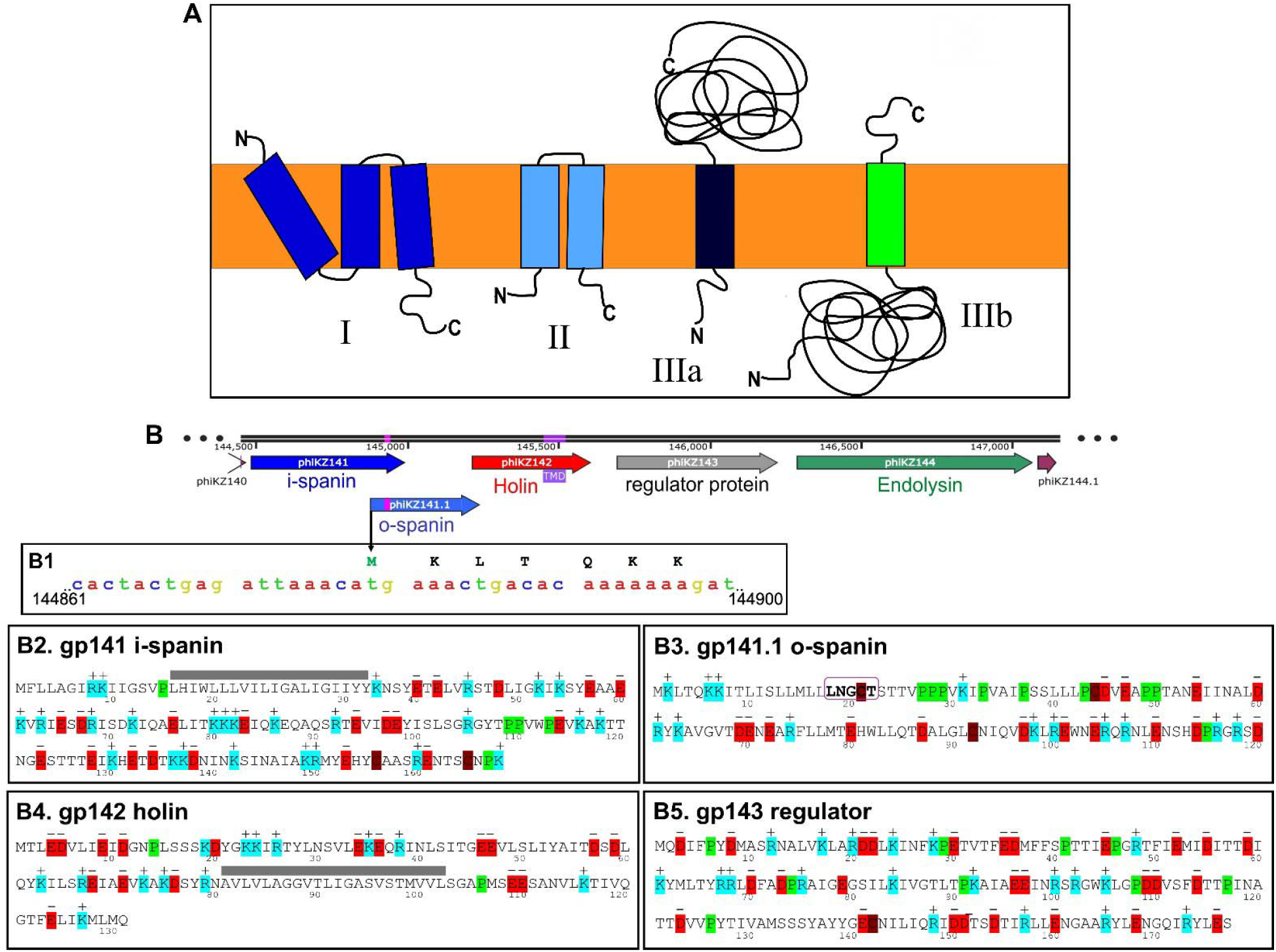
Holin topology and phiKZ lysis genes. **(A)** Topology of class I, II and III holins (Wang et al., 2000). In this study, we have grouped class III holin in phiKZ-like jumbophages as class IIIb, whereas T4-like holins are class IIIa, based on whether the large soluble domain is in the cytoplasm or periplasm. **(B)** phiKZ lysis gene cassette is drawn to scale showing the overlapping i-spanin (gp141) and o-spanin (gp141.1; pink – lipobox LNGCT boxed in bold), holin (gp142; purple – transmembrane domain), lysis regulator protein (gp143) and endolysin (gp144). **B1)** Translational start region of o-spanin. **(B2-B5)** Primary structures of all the identified lysis genes. Predicted TMDs are shown as blue bars and the Cys in lipobox of o-spanin is marked in pink.

## Materials and methods

### Bacterial strains, bacteriophages, plasmids, and growth conditions

Phages, bacterial strains, and plasmids used in this study are listed in Table 1. Bacterial strains were grown in the LB media or agar plates, supplemented with ampicillin (Amp, 100 μg/mL) or chloramphenicol (Cam, 10 μg/mL), kanamycin (Kan, 50 μg/mL) or tetracycline (Tet, 10 or 50 μg/mL) (Moussa et al., 2014; To et al., 2013). The lysogens used in this study were derived from *E. coli* K-12 MG1655 *lacI*^*Q*^ *ΔlacY* (λ *stf::cam cI*_*857*_ *bor::kan*). The λ prophages differed only in the lysis genes: RY37719 (*S*_*am*_*R*^*+*^*Rz*^*+*^*Rz1*^+^) and RY37463 (*S*^*+*^*R*^*+*^*Rz*_*am*_*Rz1*_*am*_). Bacteria were incubated at 30°C for λ *cI*_*857*_ lysogens and 37°C for non-lysogenic *E. coli* strains. MG1655 strains harboring the pBAD24 plasmid were induced by adding arabinose (final concentration of 0.4%, v/v) (Guzman et al., 1995). pQFT plasmids were induced with 1.5 mM 4-isopropylnenzoic acid (cumate) (Klotz et al., 2023). Subcultures were prepared by diluting overnight cultures inoculated from single colonies 1:250, to A_550_ ∼ 0.04 in LB with antibiotics and grown at 30°C in the case of lysogens or 37°C. Lysogens were thermally induced at A_550_ ∼ 0.2 by shifting to 42°C for 15 min and then continued growth at 37°C until lysis was observed (Park et al., 2006). For plasmid inductions with spanin genes, lysis curves were recorded with the addition of 25 mM Mg^2+^ with vigorous agitation. Wherever indicated, bacterial cultures were treated with 1 M 2,4-dinitrophenol (DNP dissolved in DMSO). Absorbance was recorded at different time intervals and plotted as a ‘lysis curve’ using GraphPad Prism software. All the data are presented as mean ± standard deviation (SD) of at least three independent experiments with error bars. For microscopic analysis, 5 µL of culture was placed on a glass slide under a coverslip. Images were acquired with an Axiocam 702 mono camera on a Zeiss Axio Observer 7 inverted microscope at 100× magnification.

**Table 1.**
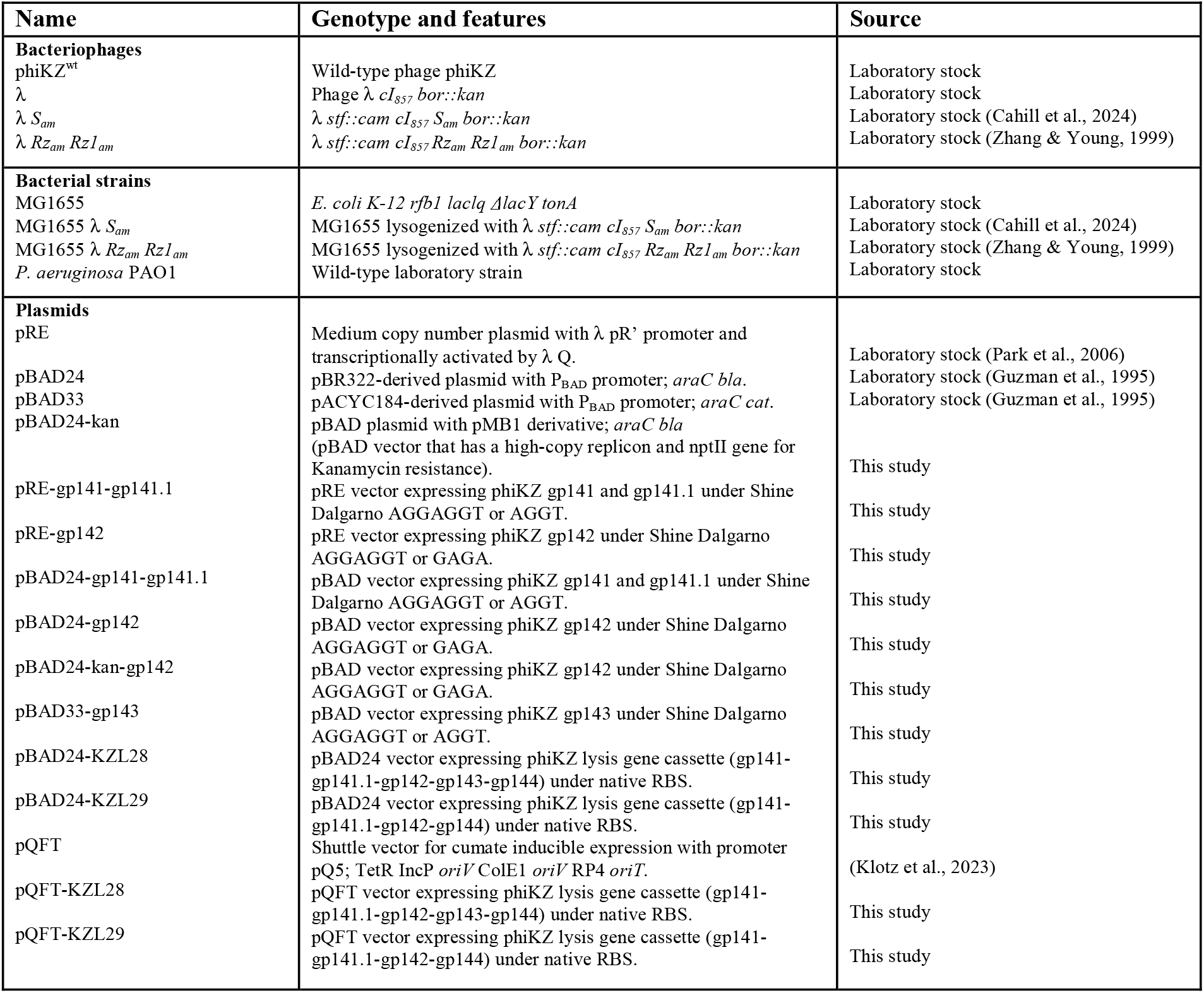
Bacteriophage, bacterial strains and plasmids used in this study.

Table 1 lists the plasmids used and constructed in this study, while table S1 provides the primers. λ lysogens were used as templates for cloning phiKZ genes. We amplified phiKZ genes *141, 141*.*1* (reannotated), *142*, and *143* from phiKZ DNA using unique primers. The genes *141, 141*.*1*, and *142* were cloned into pRE, medium copy number plasmid with λ pR’ promoter and transcriptionally activated by λ Q (Park et al., 2006). The lysis genes were cloned in pRE between its unique PstI and HindIII sites. Similarly, the genes *141, 141*.*1, 142* and *143* were cloned into pBAD, which is a pMB1 derivative with araC, bla, and a high copy replicon. All the plasmids were expressed in a temperature-sensitive λ lysogen system or MG1655. For normal cloning experiments, the Gibson assembly method was used, and the primers were made with either the native Shine-Dalgarno (S-D) sequence or the strong S-D sequence (AGGAGGT). The lysis gene cassette from nucleotide 144,479 to 147,070 (includes gp*141, 141*.*1, 142, 143*, and *144*), was cloned in pBAD for expression in MG1655, and cumate-inducible pQFT (7.5 kb) for expression in *P. aeruginosa* PAO1.

### Bioinformatics analysis

The phiKZ genome was analyzed using NCBI BLASTP and PsiBLAST for similarity searches and Artemis (Carver et al., 2012) for genome display and editing. The amino acid sequence was analyzed using DeepTMHMM (Hallgren et al., 2022), TOPCONS (Tsirigos et al., 2015), HHPred (Zimmermann et al., 2018), InterProScan (Mitchell et al., 2019), SignalIP (Teufel et al., 2022), LipoP (Juncker et al., 2003a), and Waggawagga (Simm et al., 2015) for domain and functional analysis. Full amino acid sequences were aligned using CLC Main Workbench. A multi-sequence alignment (MSA) was built against the closely related protein sequences.

Structural prediction of phage proteins was obtained in AlphaFold3 (AF3) server (Abramson et al., 2024a) (https://alphafoldserver.com). The amino acid sequences used for prediction was obtained from NCBI accession numbers, phiKZ (NC_004629.1), Noxifer (NC_041994.1), 201phi2-1 (NC_010821.1), phiPA3 (NC_028999.1) Viktualia (PV037725), Phabio (NC_062582.1) and HPP-Temi (PP968062). After the process was finished, we used pLDDT to evaluate the reliable prediction and pTM (predicted template modelling) score of > 0.5 indicates high similarity.

## Results and Discussion

### Spanin genes of phage phiKZ

Many tailed phages cluster their lysis genes in “lysis cassettes”, so our attention was drawn to genes near gene *144*, which encodes the well-studied endolysin, a transglycosylase (Fokine et al., 2008). Gene *141* was annotated as an “Rz-like spanin” in the Refseq genome (<u>NC_004629.1</u>) but this was open to question because gp141 had no detectable sequence similarity to an established i-spanin and there was no cognate o-spanin identified. We examined gene *141* for features that would be consistent with i-spanin function. The most important feature would be overlap or adjacency with the o-spanin gene, which is invariably the next gene downstream from the i-spanin (Kongari et al., 2018). Inspection of the RefSeq phiKZ genome (NC_004629.1) revealed that upstream of the start codon of the adjacent CDS, gene *141*.*1*, there were 5 ATG or GTG start codons in frame, only one of which had a Shine-Dalgarno sequence appropriately placed for its start codon (Fig.1B). Using this start codon, this version of gene *141*.*1* spans 120 codons and, upon analysis by LipoP (Juncker et al., 2003b) was predicted to encode a lipoprotein destined for the inner leaflet of the OM (lipobox LNGCT; (Braun & Wu, 1994)) (Fig.1B). Thus, we designated it as the o-spanin gene for phiKZ, strongly indicating that gene *141* was the i-spanin gene. The periplasmic domain of i-spanins is dominated by alpha-helical structure and the presence of a coiled-coil structure, as garnered from biochemical and structural experiments with the lambda Rz protein (J. Berry et al., 2010a, 2010b; Cahill, Rajaure, O’Leary, et al., 2017) and extensive computational analysis of a large and very diverse set of i-spanins (Kongari et al., 2018). Gp141 is predicted to have these features (Fig.2A). To test these assignments, the *141/141*.*1* overlapping gene pair was cloned into the trans-activation vector pRE (Park et al., 2006) and transformed into a lambda lysogen carrying nonsense mutations in the *Rz* i-spanin and *Rz1* o-spanin genes. In this construct, the plasmid-borne spanin genes are controlled by the Q late gene anti-terminator encoded on the prophage (Grayhack & Roberts, 1982) and are thus expressed (transactivated) at the appropriate time after lysogenic induction (Park et al., 2006). Under these conditions, the *141/141*.*1* gene pair fully complemented the lysis defect of λ*Rz*_*am*_*Rz1*_*am*_ prophage, where the non-complementation terminal phenotype after induction is spherical cells bounded by the intact OM (Cahill, Rajaure, Holt, et al., 2017) (Fig.2B).

**Figure 2.**
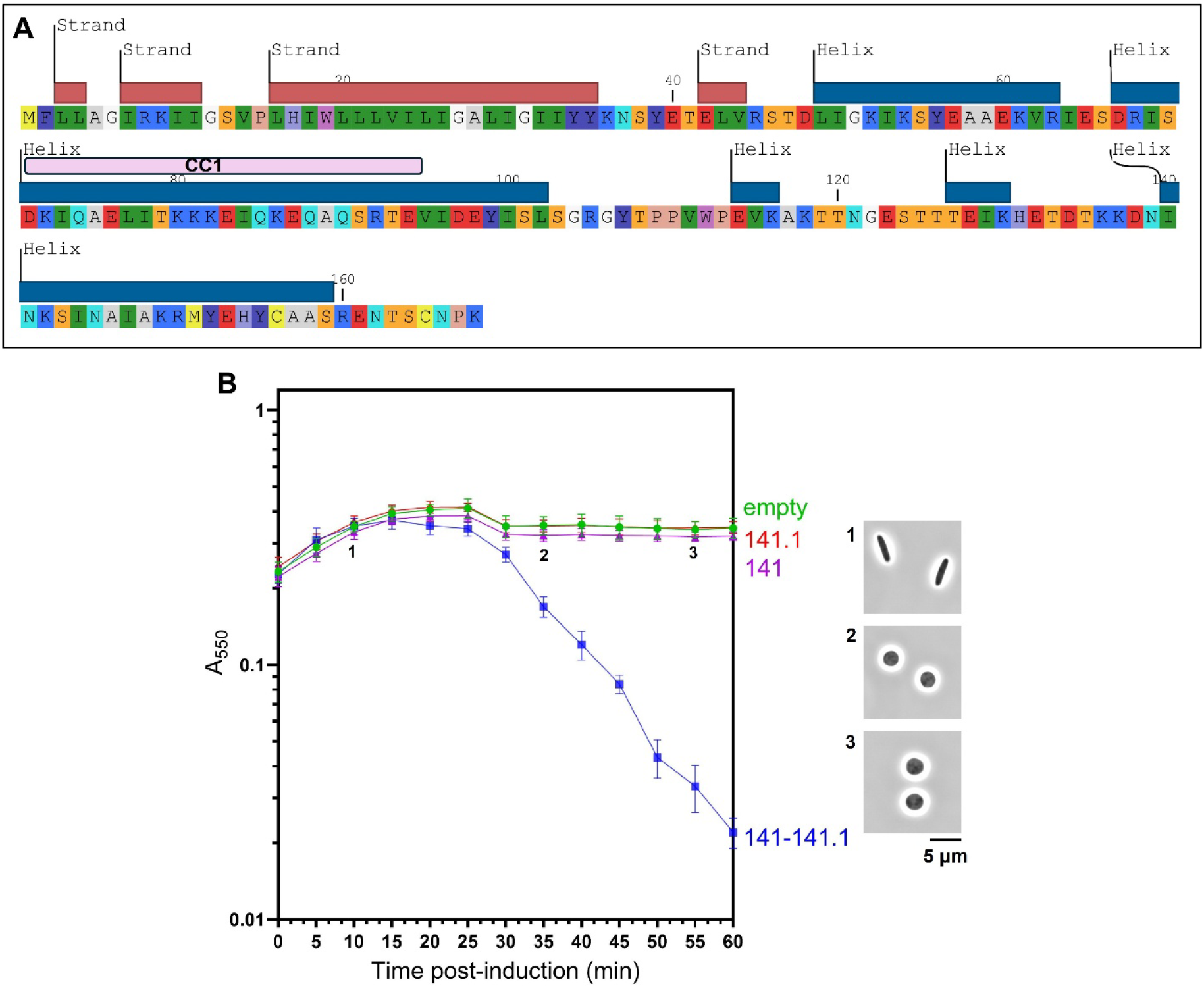
phiKZ spanin phenotypes. **(A)** Predicted secondary structure of i-spanin, gp141. Blue and red rectangles indicate predicted alpha helical and beta strand/ sheet domains respectively. Coiled-coil domain CC1 is highlighted in pink rectangle. (**B)** gp141 and gp141.1, i-spanin and o-spanin of phiKZ, in pRE vector were thermically induced in *E. coli* RY37463 (MG1655 *lacI*^*q*^*ΔlacY* carrying the prophage λ *cI857 S*^*+*^*R*^*+*^*Rz*_*am*_*Rz1*_*am*_, *stf::cam cI*_*857*_ *bor::kan*) with the addition of 25 mM Mg^2+^ in the following combinations: empty pRE vector (green), pRE-gp141 (purple), pRE-gp141.1 (red) and pRE-gp141-gp141.1 (blue). At 10, 35 and 60 min (marked as 1, 2, and 3) pRE-gp141 and pRE-gp141.1 samples were observed at 100× magnification to observe the cell morphology. The data represents the mean ± standard deviation of three independent experiments.

### The phiKZ holin

The unambiguous identification of the spanin genes left genes *142* and *143* to be considered for the missing holin assignment, if a lysis cassette is indeed present. All holins have at least one TMD (Cahill & Young, 2019), so gp*143* can be ruled out (Fig.1A; Supplementary data S1). Gp*142* has a single TMD predicted by TMHMM and other membrane helix propensity programs (see Methods). We hypothesized that gp142 was a class III holin, a class founded by the well-studied gp*t* of the canonical myophage T4 (Fig.1A). Whereas other established classes of holins, classes I (3 TMDs, N-out, C-in) and II (2 TMDs, N-in, C-out) are extremely diverse and found in all types of *Caudoviricetes* (Young, 2002), all the members of class III (1 TMD, N-in, C-out) were close homologs of gp*t*, including *hol* of phage T5 (Chernyshov et al., 2024), and found only in the phages broadly characterized as T4-like and T5-like phages (Summer et al., 2007). Gp*t* has been extensively studied at the genetic, physiological and structural levels (Moussa et al., 2014; Ramanculov & Young, 2001b, 2001a; Tran et al., 2005), especially its large periplasmic domain (Fig.1). Based on these studies, the current model for gp*t*-mediated lysis involves hole formation requiring the single TMD and negatively regulated by interaction with a periplasmic antiholin, gp*rI* and a cytoplasmic anti-holin, gp*rIII* (Fig.1A). Gp*142* has the same gross topology as gp*t*, with a single N-in, C-out TMD, but the sizes of the soluble domains are reversed, with a large cytoplasmic and a small periplasmic domain in gp*142*. Although extremely diverse, holins have several universal functional features, the most central of which is a temporally regulated lethality, as briefly noted above (Bavda & Jain, 2020; Ramanculov & Young, 2001b; Wang et al., 2000). We cloned gene *142* into the transactivation plasmid and tested for its ability to complement the lytic function of the classical lambda holin, *S*_*105*_ (Smith et al., 1998). The results unambiguously established that gp*142* has holin function capable of allowing the escape of a soluble globular endolysin, the R transglycosylase of phage lambda (Bläsi et al., 1999) (Fig.3A). Moreover, gp*142* exhibited the definitive feature of being sensitive to premature triggering by collapse of the proton motive force (PMF) by addition of the uncoupler dinitrophenol (Raab et al., 1986; Reader & Siminovitch, 1971) (Fig.3B). Moreover, premature triggering is instantaneous and very rapid unless the energy poison is added too early in the latent period, in which case lysis is more gradual, presumably reflecting an insufficient level of holin expression (Reader & Siminovitch, 1971). This behavior is also observed with gene *142*. Taken together, these results establish gp*142* as the holin of phiKZ.

**Figure 3.**
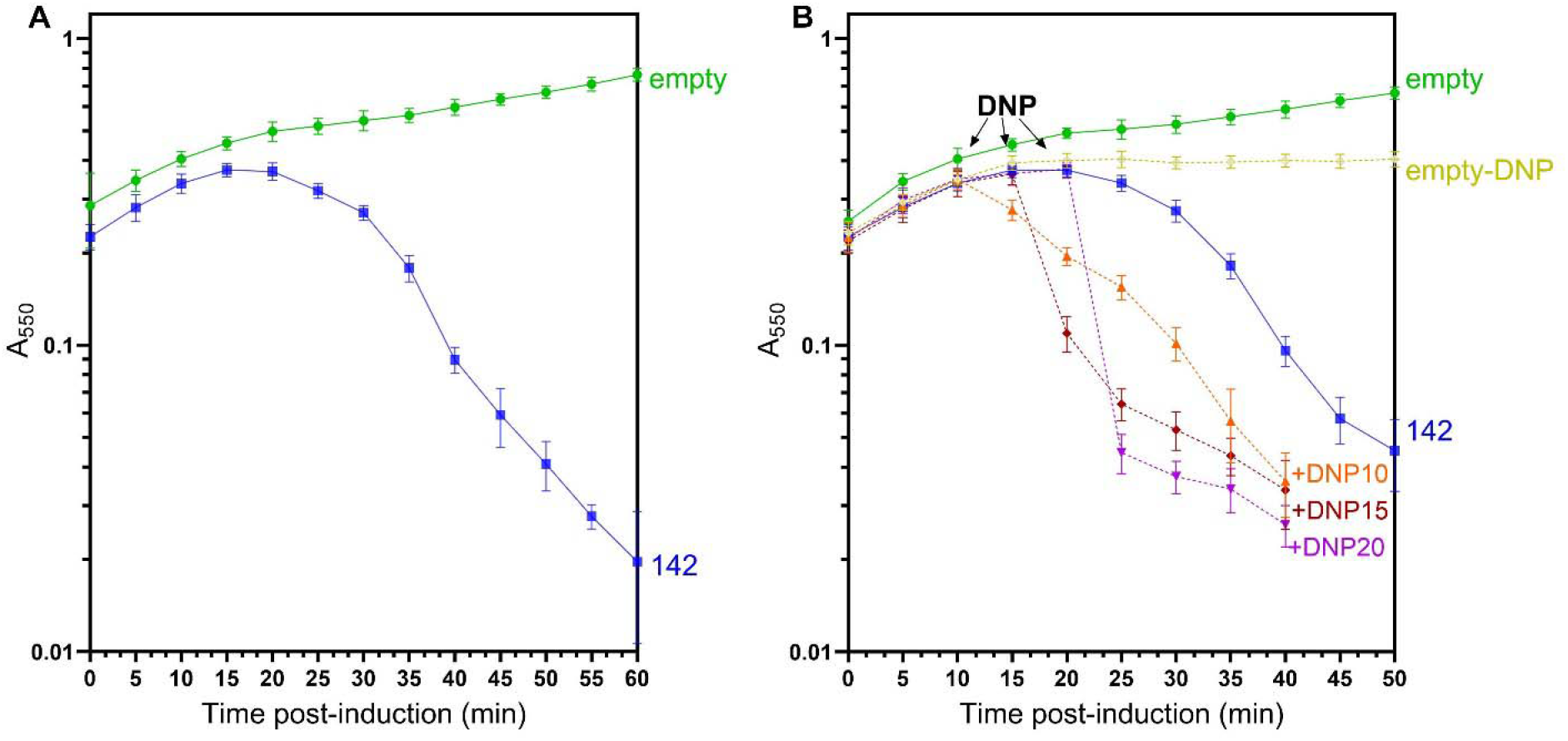
phiKZ holin gene lysis. **(A)** gp142, holin of phiKZ, in pRE vector was induced with arabinose in *E. coli* RY37463 (MG1655 *lacI*^*q*^*ΔlacY* carrying the prophage λ *cI857 S*^*+*^*R*^*+*^*Rz*_*am*_*Rz1*_*am*_, *stf::cam cI*_*857*_ *bor::kan*) and lysogens were thermally induced at 42°C in the following combinations: empty pRE vector (green), and pRE-gp142 (blue). (**B)** MG1655 lysogens containing pBAD24-gp142 (blue) were thermally induced at 42°C and shifted to 37°C to induce with arabinose. At 10, 15 and 20 min, cells were treated with 1mM DNP (dashed lines). The data represents the mean ± standard deviation of three independent experiments.

### Gp143: lysis regulator

The assignments to genes *141, 141*.*1, 142* and *144* account for all four standard types of MGL proteins. As a true lysis cassette and considering that MGL systems are often functional when expressed in heterologous backgrounds (Briers et al., 2011; Cahill et al., 2024), a plasmid with all four genes should cause lysis when expressed in *E. coli*. An arabinose-inducible plasmid carrying five genes, the putative four gene lysis cassette plus the intervening gene *143*, in their native translational context, was constructed and used to transform *E. coli*. Induction with arabinose of mid-logarithmic culture of the transformant resulted in a sharply defined lysis profile at ∼45 min after induction (Fig.4A,B). To see if gp*143* was involved, the experiment was repeated with an isogenic plasmid where gene *143* was deleted. In this case lysis was delayed by ∼15 min but retained its saltatory character. This partial defect was also found in the *Pseudomonas* context using plasmid constructs based on the pQFT cumate-inducible vector (Fig.4C). This finding could be due to reduced expression of the holin, although gene *143* being downstream of the holin gene makes this less likely. Accordingly, the *142* holin gene and the *143* gene were cloned into compatible ara-inducible plasmids, transformed into *E. coli* and tested for the inducible lethality that is the hallmark of holin genes (Fig.4D). The results showed that the expression of gene *143* in trans caused triggering ∼10 min earlier than with the holin alone. It is unlikely this is due to increased translation of the holin mRNA, since in these constructs the translational initiation sequences are derived from the vector. In both cases the induced lethality is observed in the sudden cessation of logarithmic growth; in addition, there is a gradual and limited decrease in culture density, due to vacuolization that reflects the permeabilization of the cytoplasmic membrane, the first step in MGL lysis, in the absence of endolysin and spanin function (Fig.1D) (Cahill et al., 2024). The simplest explanation is that gp143 interacts with the holin to regulate hole-formation, as has been found in three cases where a protein has been shown to affect holin timing (Barenboim et al., 1999; Bläsi & Young, 1996; Ramanculov & Young, 2001a). Since gp143 has no predicted signal sequence or TMD and is thus predicted to be a cytoplasmic protein, the unusually long N-terminal cytoplasmic domain of gp142 holin was the most attractive candidate for a regulatory interaction site. Accordingly, we generated an isogenic construct carrying a deletion allele of the holin gene, *142*_*Δ(2-60)*,_ encoding a holin protein lacking most of the N-terminal cytoplasmic domain (Fig.5A). This truncation construct was tested in *E. coli* and found to be fully functional in terms of inducible lethality (Fig.5B), although triggering ∼15 min later than the wt. Strikingly the expression of gp143 in trans blocked the triggering of the truncated holin, instead of accelerating triggering as observed for the wt holin allele. Although more studies will be needed to illuminate the molecular mechanism involved, together these *in vivo* results indicate that gp143 is a holin-specific lysis regulator depending on interaction with the N-terminal cytoplasmic domain of the gp142 holin. Thus, every one of the five adjacent cistrons beginning with *141* is involved in MGL lysis and thus can unambiguously be termed a lysis cassette.

**Figure 4.**
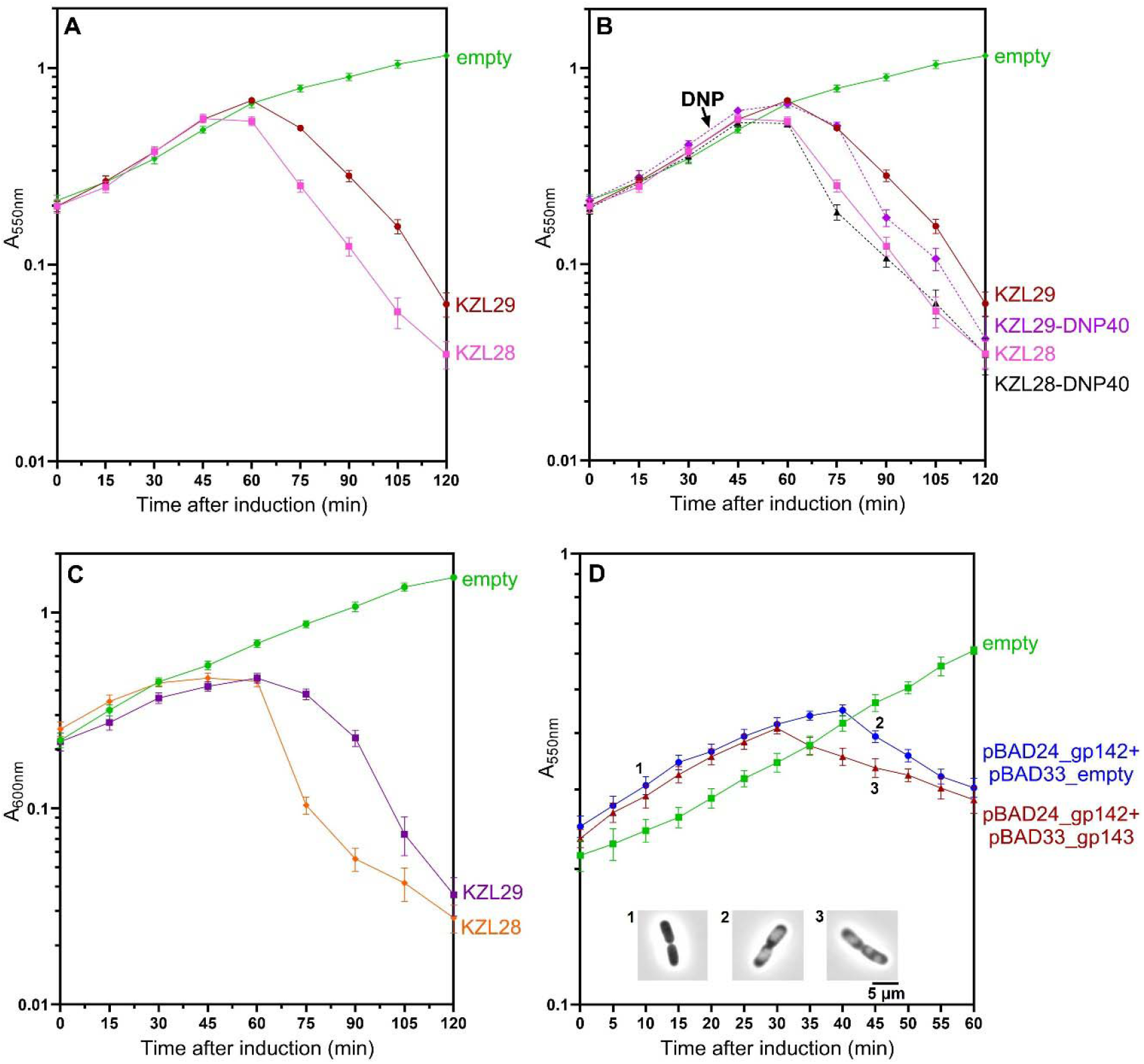
phiKZ lysis gene cassette with a regulator gene. **(A)** The lysis gene cassette (gp141, 141.1, 142, *143, 144) in pBAD vector were induced with arabinose in MG1655 cells grown in the presence of 25 mM Mg^2+^ in the following combinations: empty pBAD24 vector (green), pBAD24-KZL28 (gp141-141.1-142-143-144) (pink) and pBAD24-KZL29 (gp141-141.1-142-144) (maroon). (**B)** MG1655 containing pBAD24-lysis gene cassette were induced with arabinose. At 40 min, cells were treated with 1mM DNP (dashed lines). (**C**) *P. aeruginosa* PAO1 containing pQFT-lysis gene cassette were induced with 1.5 mM cumate in the following combinations: empty pQFT vector (green), pQFT-KZL28 (orange) and pBAD24-KZL29 (purple). (**D)** Co-expression of pBAD24-gp142 and pBAD33-gp143 in MG1655 cells and were induced with arabinose in the following combinations: empty pBAD24+pBAD33 vectors, pBAD24-gp142+ pBAD33-empty (green), pBAD24-gp142+ pBAD33-gp143 (maroon). At 10 and 45 min (marked as 1, 2, and 3) pBAD24-gp142+pBAD33-empty and pBAD24-gp142+ pBAD33-gp143 samples were observed at 100× magnification to observe the cell morphology and vacuolization was observed in the presence of gp142 holin. The data represents the mean ± standard deviation of three independent experiments.

**Figure 5.**
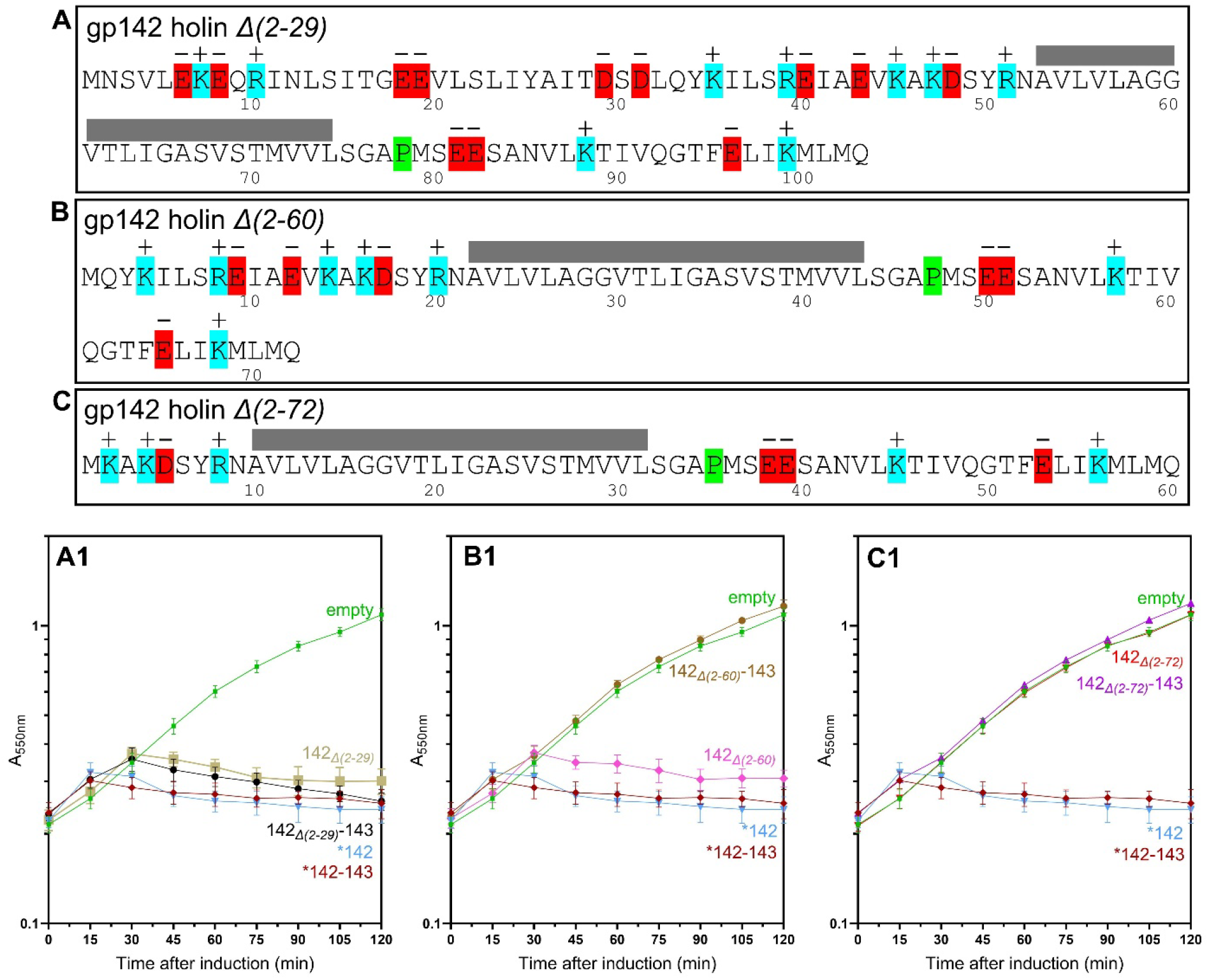
Functional analysis of phiKZ holin mutant alleles. **(A, B, C)** Primary structures of three holin mutants, gp142_*Δ(2-29)*_, gp142_*Δ(2-60)*_, and gp142_*Δ(2-72)*_. Transmembrane domain is marked in gray box. **(A1, B1, C1)** The mutant holin alleles with and without gp143 in pBAD24-kan vector were induced with arabinose in MG1655 cells in the following combinations: **(A1)** empty pBAD24 vector (green), pBAD24-gp142 (light blue) and pBAD24-gp142+143 (maroon), pBAD24-gp142_*Δ(2-29)*_ (gold), and pBAD24- gp142_*Δ(2-29)*_+143 (black). **(B1)** pBAD24-gp142_*Δ(2-60)*_ (pink), and pBAD24-gp142_*Δ(2-60)*_+143 (brown). **(C1)** pBAD24-gp142_*Δ(2-72)*_ (red), and pBAD24-gp142_*Δ(2-72)*_+143 (purple). The data represents the mean ± standard deviation of three independent experiments.

### Structural modelling of phiKZ lysis cassette protein

Current artificial intelligence software for structure prediction is problematic for use on membrane proteins (Abramson et al., 2024b; Agarwal & McShan, 2024; Niazi, 2025). Only the lysis regulator gp143 is a native of the cytosol and could be expected to generate a high confidence prediction using Alpha-fold (AF), as it did, generating a high-confidence structure with two globular domains (Fig.6A; Fig.S1). Gp142, as a predicted class III holin, has a single-pass, N-in, C-out TMD, making the full-length molecule a poor candidate for AF analysis. Accordingly, AF generated a high confidence structure for the soluble N-terminal domain (residues 1-74), although the positioning relative to the TMD is less certain. Accordingly, we used AF to predict the structure of a heterodimer of the predicted N-terminal cytoplasmic domain of the gp142 holin and the complete gp143. This resulted in a high confidence prediction, in which the first globular domain of gp143 binds to an alpha-helical bundle of gp142 through a network of hydrogen bonds (Fig.6B). This structure is largely conserved among phiKZ-like phages (Fig.S2, S3 & S4). Further genetic, biochemical and structural efforts will be needed to validate the structure and determine what role the putative heterodimer plays in regulation of lysis timing. Interestingly, liquid infection assays using low and high multiplicity of infection reveal that increasing MOI actually retards lysis (Fig.S5B). Moreover, bulk culture infections show gradual lysis extending over more than 2 h, whereas single step growth assay indicates a lysis time of 30-40 min (Fig.S5A). These findings suggest a form of lysis inhibition (LIN) (V. Krylov et al., 2021; Pleteneva et al., 2010). The classic LIN system of T4 offers parallels with phiKZ, with the major holin regulator, RI, binding to a globular domain of a class III holin. However, the binding occurs in the periplasm in the case of T4. Moreover, in the absence of superinfection in phiKZ infections, the effect of the binding appears to be positive, rather than negative (Fig.S6). In any case, given phiKZ’s status as the prototype of the nucleus-forming phages, the lysis and putative LIN system merit further study.

**Figure 6.**
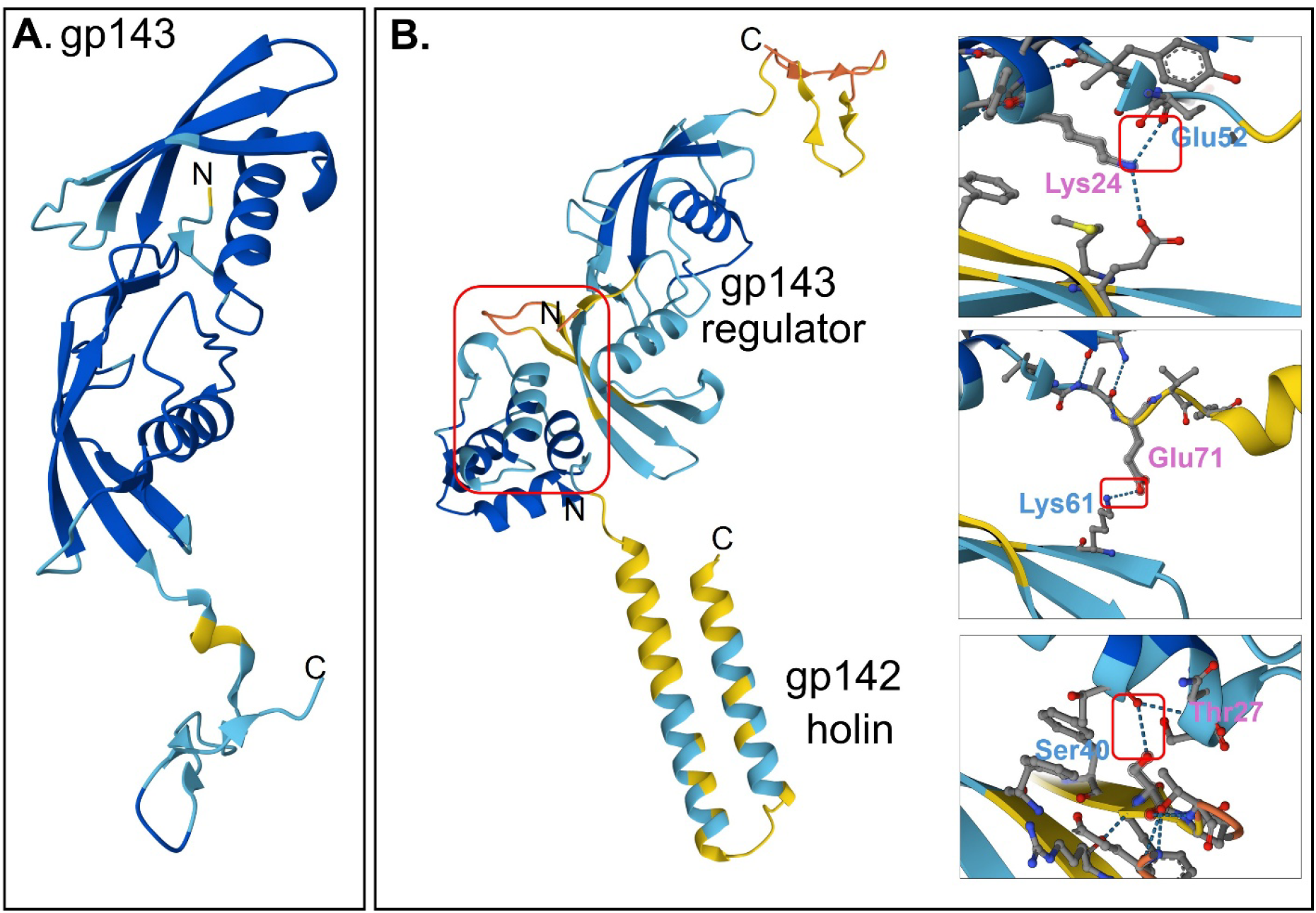
AlphaFold3 model of phiKZ gp143 and its interaction with gp142. **(A)** Predicted structure of gp143. **(B)** Interaction between N-terminal domain of gp142 (first 80 residues are predicted to be in the cytoplasm) and gp143 (boxed), where N-terminal gp142 residues Lys24, Glu71 and Thr27, and gp143 residues Glu52, Lys61 and Ser40 are involved in protein-protein interaction. See figure S1 for PAE plots. 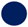 pLDDT > 90 (very high similarity), 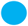 pLDDT of 70-90 (high similarity), 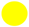 pLDDT of 50-70 (low similarity), 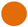 pLDDT < 50 (very low similarity).

## Acknowledgment

The authors would like to thank the funding supported by the National Institute of General Medical Sciences, grant no. R35GM136396 to R.Y. and by the Center for Phage Technology, which is jointly supported by Texas A&M AgriLife Research.

## Authorship contribution

Conceptualization, P.M., J.W., R.Y.; Data Curation, P.M., J.W.; Formal Analysis, P.M., G.G., K.S., J.W.; Experimental works, P.M., J.W., G.G., K.S.; Investigation, P.M., J.W.; Methodology, P.M., J.W., R.Y.; Resources, P.M., R.Y.; Supervision, R.Y.; Writing – Original Draft, P.M., R.Y.; Writing – Review & Editing, P.M., R.Y.; Validation, P.M., R.Y.; Visualization, P.M.; Project Administration, P.M., R.Y.; Funding Acquisition, R.Y.

## Supplementary Information

**Supplementary table S1:** Oligonucleotides used in this study.

## Supplementary figures

Figure S1: Comparison of predicted alignment error (PAE) plots of phiKZ from Alphafold3.

Figure S2: Alignment of the similarity and domain composition of (A) holin and (B) phylogenetic tree of holins among all the phiKZ-like jumbo phages.

Figure S3: Alignment of the similarity among (A) lysis regulator protein and (B) phylogenetic tree of lysis regulator proteins among all the phiKZ-like jumbo phages.

Figure S4: AlphaFold3 model to show the interaction between holin and lysis regulator in three phiKZ-like phages.

Figure S5: *Pseudomonas* phage phiKZ growth kinetics.

Figure S6: Functional analysis of phiKZ holin mutant alleles.

**Supplementary data 1:** List of all the phiKZ genes with transmembrane domain (TMD). The results or output was generated using TMHMM webserver.

