## Supplemental Table for "The Lysis Cassette of Jumbophage phiKZ"

### Supplementary material:

#### Supplemental table:

#### Table S1: Oligonucleotides used in this study.

| Primer name | Sequence (5’-3’) | Purpose |
|---|---|---|
| KZ141+141.1 F | atgttcctattagcaggtatacga | Amplify KZ141,141.1 |
| KZ141+141.1 R | tcaatcagaacgtcctctagggtc | Amplify KZ141,141.1 |
| KZ142 F | atgaccctagaggacgttctgattga | Amplify KZ142 |
| KZ142 R | ttattgcataagcatctttattaattc | Amplify KZ142 |
| pBAD24 PCR F | ggctgttttggcggatgag | Linearize pBAD24 |
| pBAD24 PCR R | ggtgaattcctcctgctag | Linearize pBAD24 |
| araC-pBAD Gib F | tgctggccttttgctcacatgttctttcctgcgttatcccatcgatgcataatgtgcctgtca | araC-pBAD24 |
| araC-pBAD Gib R | gaaaacctctgacacatgcagctcccggagacggtcacagagagtttgtagaaacgcaaaaaggc | araC-pBAD24 |
| pBAD-kan bla F | actcttcctttttcaatattattgaagcatttatcagg | pBAD without bla |
| pBAD-kan bla R | ctgtcagaccaagtttactcatatatactttagattgatttaaaac | pBAD without bla |
| nptII Gib pBAD-kan F | aatcaatctaaagtatatatgagtaaacttggtctgacagtcagaagaactcgtcaagaaggcga | nptII for pBAD-kan |
| nptII Gib pBAD-kan R | accctgataaatgcttcaataatattgaaaaaggaagagtatgattgaacaagatggattgcacg | nptII for pBAD-kan |
| pBAD-kan len F | gacctgcaggcatgcaagc | Linearize pBAD-kan |
| pBAD-kan len R | gaattcctcctgctagcccaaaaa | Linearize pBAD-kan |
| KZ141, 141.1 Gib F | ctccatacccgtttttttgggctagcaggaggaattcaccatgttcctattagcaggtatacgaaaaataattggtag | clone gp141, 141.1 in pBAD-kan |
| KZ141,141.1 Gib R | gtatcaggctgaaaatcttctctcatccgccaaaacagcctcaatcagaacgtcctctagggtc | clone gp141, 141.1 in pBAD-kan |
| KZ142 Gib F | ctccatacccgtttttttgggctagcaggaggaattcaccatgaccctagaggacgttctgattga | clone gp142 in pBAD-kan |
| KZ142 Gib R | gtatcaggctgaaaatcttctctcatccgccaaaacagccttattgcataagcatctttattaattcaaaggtaccttgcacaatagtc | clone gp142 in pBAD-kan |
| pBAD24-Lys Gib F | ctccatacccgtttttttgggctagcaggaggaattcaccatgttcctattagcaggtatacgaaaaataattggtag | clone lysis cassette in pBAD24 |
| pBAD24-Lys Gib R | gtatcaggctgaaaatcttctctcatccgccaaaacagccttattttctatgtgctgcaactttaccatccattaagttataaa | clone lysis cassette in pBAD24 |
| pBAD24-Lys KLD F | gtaataccctcaatgaacctaacccagg | Lysis cassette without gp143-pBAD |
| pBAD24-Lys KLD R | gtcaaacctctatagtggtaaaatgtttaatccac | Lysis cassette without gp143-pBAD |
| pQFT V len F | ggatccggtgaagtgacccg | Linearize pQFT vector for Gibson |
| pQFT V len R | cggaagcttcctctactagttacaaacag | Linearize pQFT vector for Gibson |
| pQFT-Lys Gib F | gacaatctggtctgtttgtaactagtagaggaagcttccgatgttcctattagcaggtatacgaaaaataattggtag | clone lysis cassette in pQFT |
| pQFT-Lys Gib R | tcccctggcctttttttttgcgggtcacttcaccggatccttattttctatgtgctgcaactttaccatccattaagttataaa | clone lysis cassette in pQFT |
| pQFT-Lys KLDGib F | gacaatctggtctgtttgtaactagtagaggaagcttccgatgttcctattagcaggtatacgaaaaataattgg | Lysis cassette without gp143 |
| pQFT-Lys KLD Gib R | tcccctggcctttttttttgcgggtcacttcaccggatccttattttctatgtgctgcaactttacca | Lysis cassette without gp143 |
