## Supplemental Figures for "The Lysis Cassette of Jumbophage phiKZ"

### Supplementary material:

#### Supplemental figures:

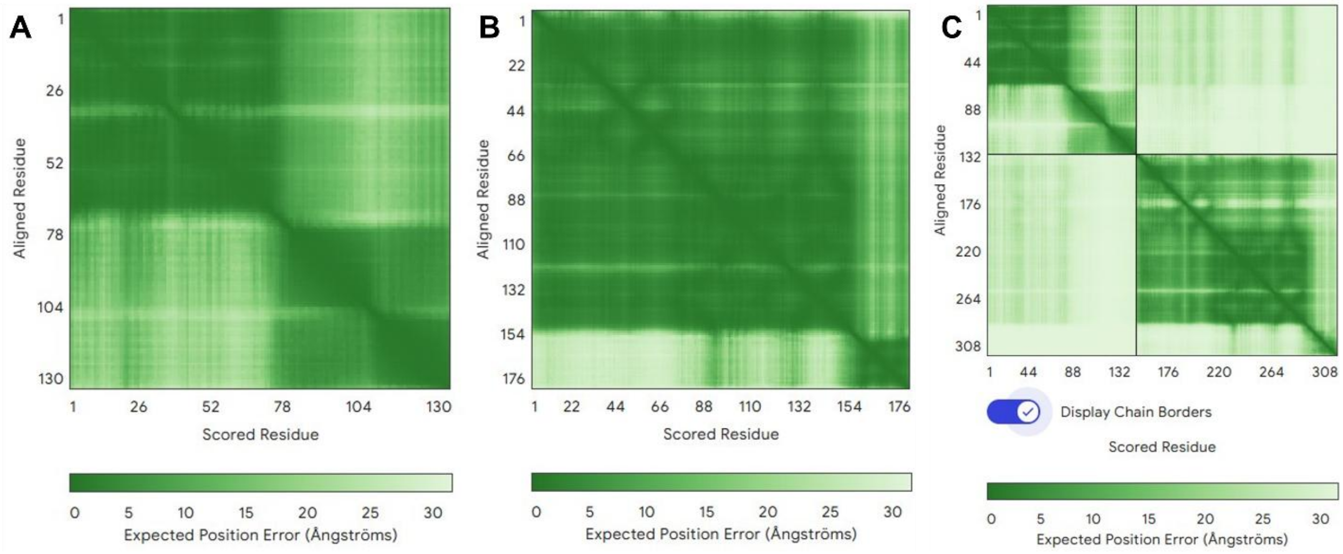


**Figure S1: Comparison of predicted alignment error (PAE) plots of phiKZ.** (A) gp142 holin, (B) gp143 lysis regulator, and (C) interaction of gp142+143. The results are obtained from AlphaFold3 server.

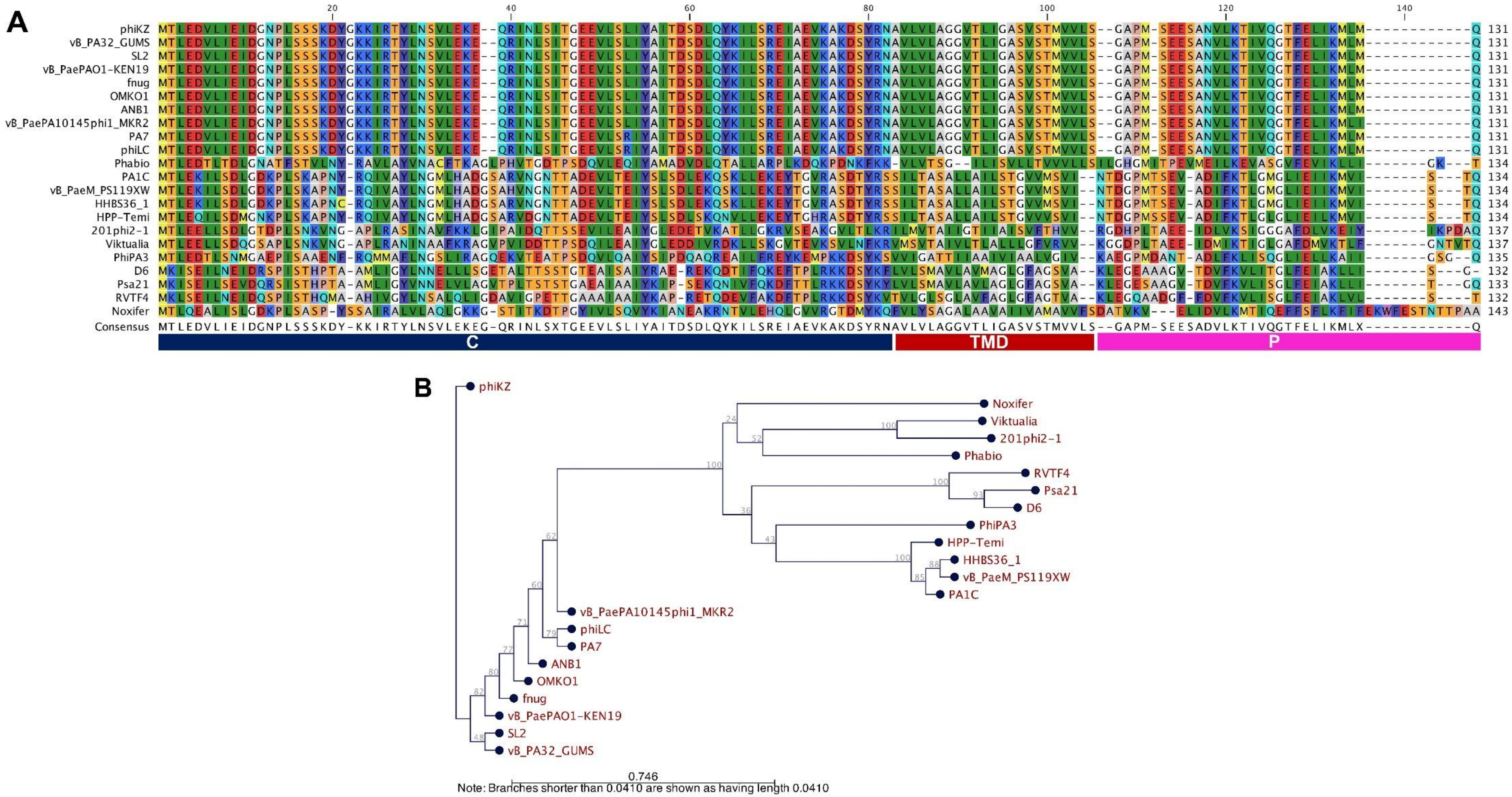


**Figure S2: Alignment of the similarity and domain composition of (A) holin and (B) phylogenetic tree of holins among all the phiKZ-like jumbo phages.** Holin is aligned with their respective TMDs. Protein sequences are aligned and labeled according to phage name. The phylogenetic tree was constructed using CLC workbench with Neighbor-joining (NJ) method and JTT matrix-based model with the bootstrap replicate of 1000. The percentage of trees on which the associated taxa clustered together is shown above the branches. This analysis involved 22 amino acid sequences.

A

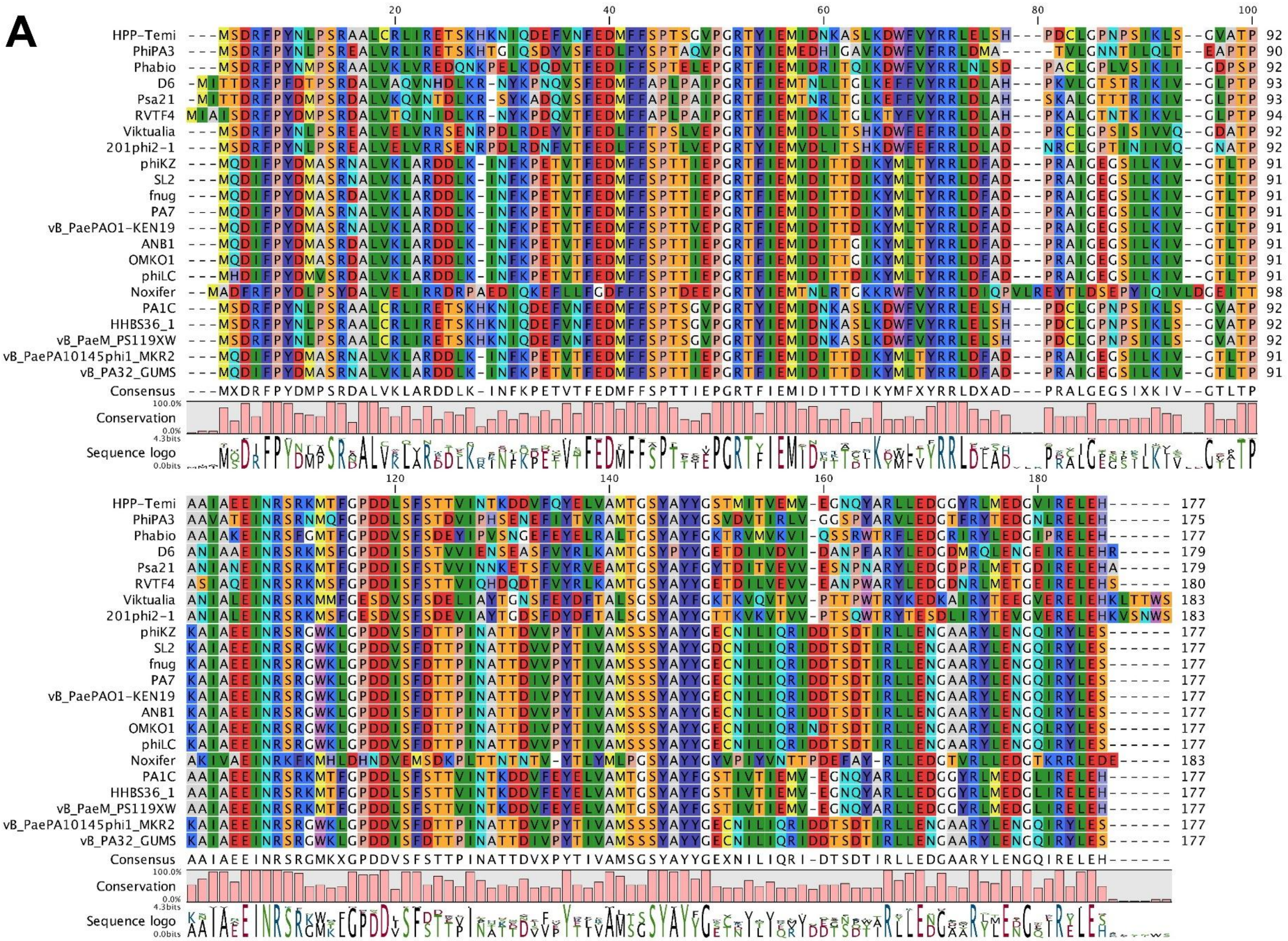
HPP-Temi
PhiPA3
Phabio
D6
Psa21
RVTF4
Viktualia
201phi2-1
phiKZ
SL2
fnug
PA7
vB_PaePAO1-KEN19
ANB1
OMKO1
phiLC
Noxifer
PA1C
HHBS36_1
vB_PaeM_PS119XW
vB_PaePA10145phi1_MKR2
vB_PA32_GUMS
Consensus
Conservation
Sequence logo

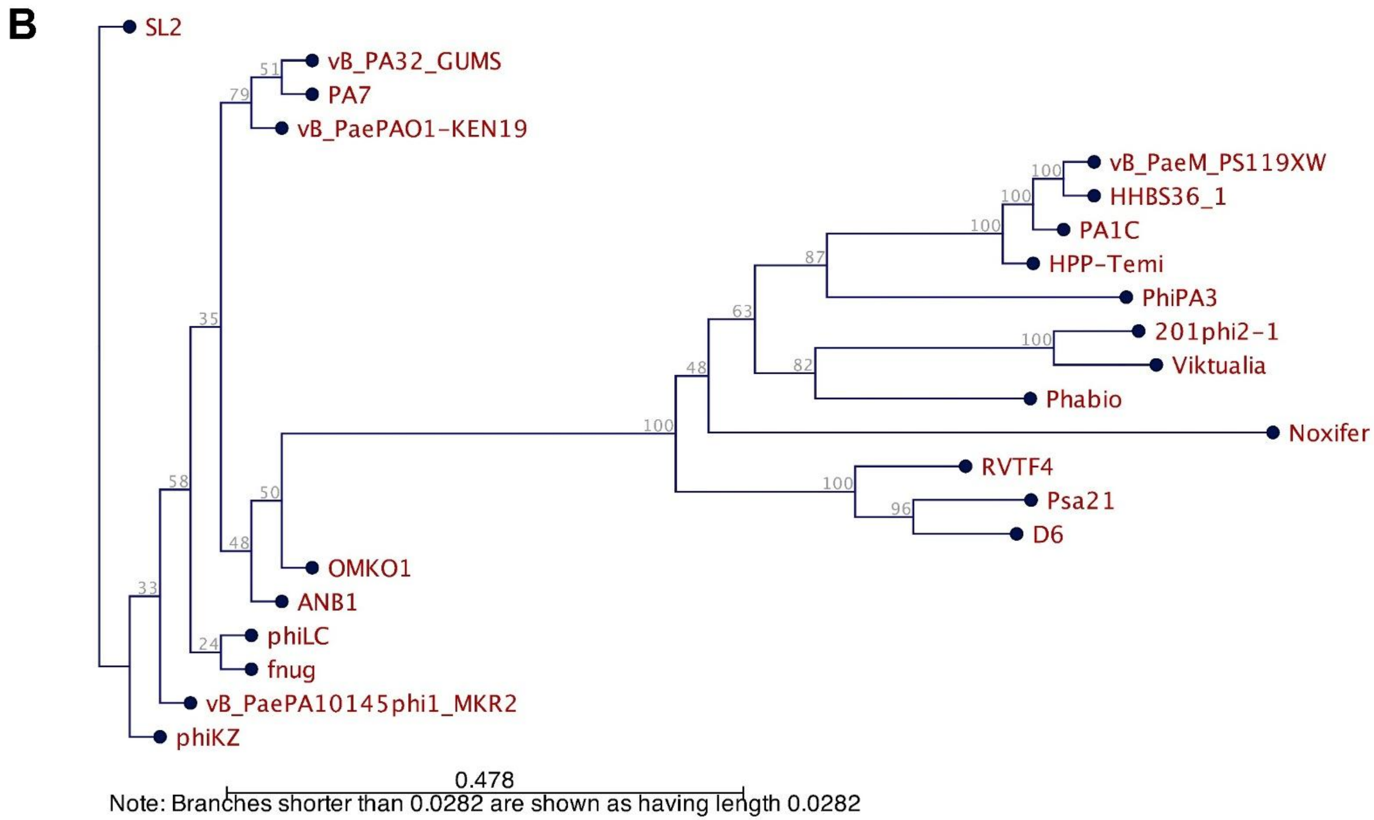


**Figure S3: Alignment of the similarity among (A) lysis regulator protein and (B) phylogenetic tree of lysis regulator proteins among all the phiKZ-like jumbo phages.** Protein sequences are aligned and labeled according to phage name. The phylogenetic tree was constructed using CLC workbench with Neighbor-joining (NJ) method and JTT matrix-based model with the bootstrap replicate of 1000. The percentage of trees on which the associated taxa clustered together is shown above the branches. This analysis involved 22 amino acid sequences.

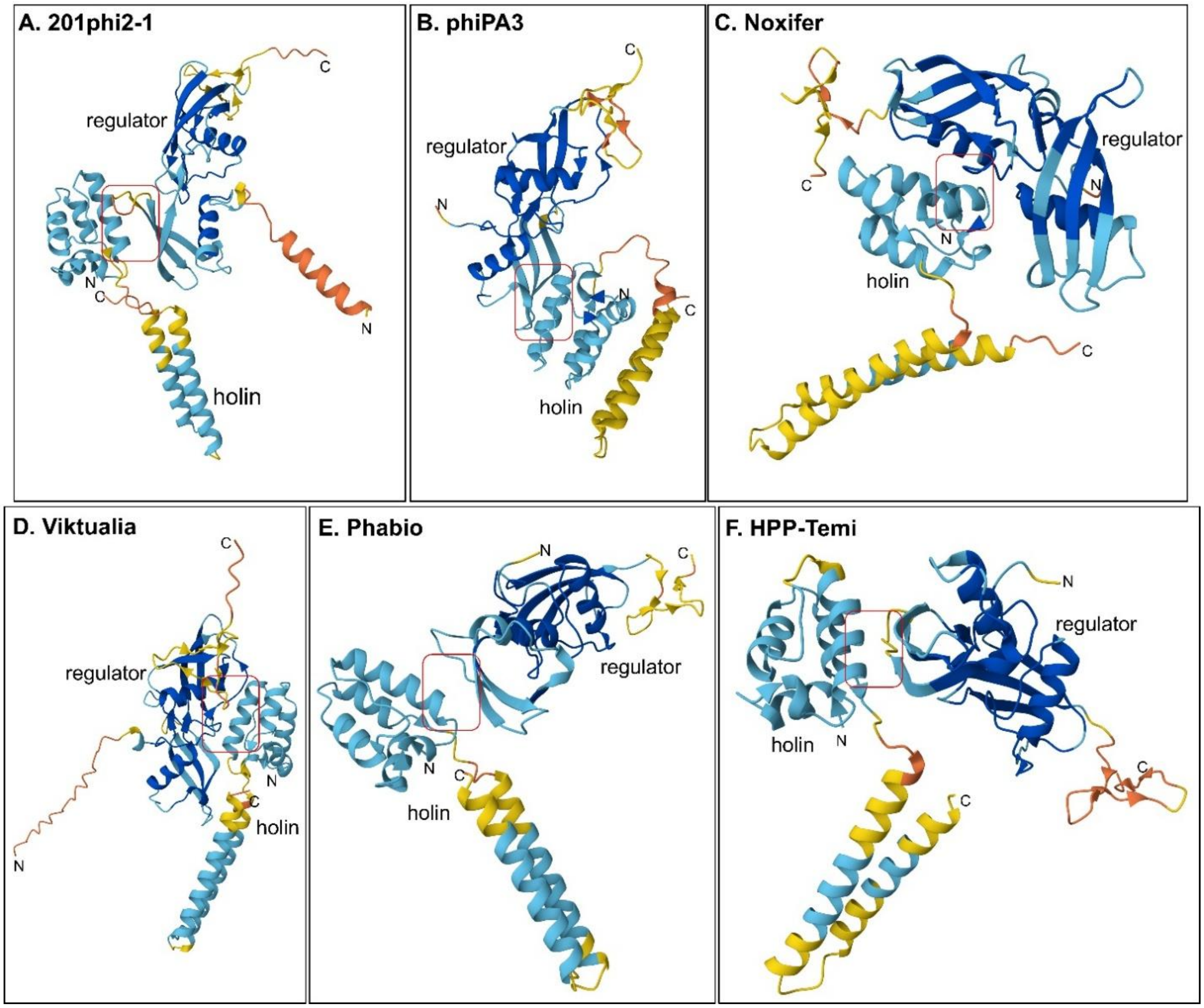


**Figure S4: AlphaFold3 model to show the interaction between holin and lysis regulator in six phiKZ-like phages.** (**A**) 201phi2-1 (NC_010821.1) [holin: ST201phi2-1p227 and regulator: ST201phi2-1p228], interaction is between N-terminal domain of holin and regulator, i.e., Ala26-Ser64 and Arg25-Glu69. (**B**) phiPA3 (NC_028999.1) [holin: AVT69_gp159 and regulator: AVT69_gp160], interaction is between N-terminal domain of holin and regulator, i.e., Arg22-Glu13 and Arg68-Glu55. (**C**) Noxifer (NC_041994.1) [holin: NOXIFER_142 and regulator: NOXIFER_141], interaction is between N-terminal domain of holin and regulator, i.e., Ile8-Asn106 and Ser9-His112. (**D**) Viktualia (PV037725) [holin: APQOYWFA_CDS0435 and regulator: APQOYWFA_CDS0436], interaction is between N-terminal domain of holin and regulator, i.e., Asn27-Glu132, Lys68-Glu132, Arg34-Ser136 and Asn20-Try160. (**E**) Phabio (NC_062582.1) [holin: MZD05_gp215 and regulator: MZD05_gp216], interaction is between N-terminal domain of holin and regulator, i.e., Arg22-Asp63. (**F**) HPP-Temi (PP968062) [holin: HSP1_CDS0261 and regulator: HSP1_CDS0260], interaction is between N-terminal domain of holin and regulator, i.e., Tyr27-Ser41 and Gln23-Phe39. ● pLDDT > 90 (very high similarity), ● pLDDT of 70-90 (high similarity), ● pLDDT of 50-70 (low similarity), ● pLDDT < 50 (very low similarity).

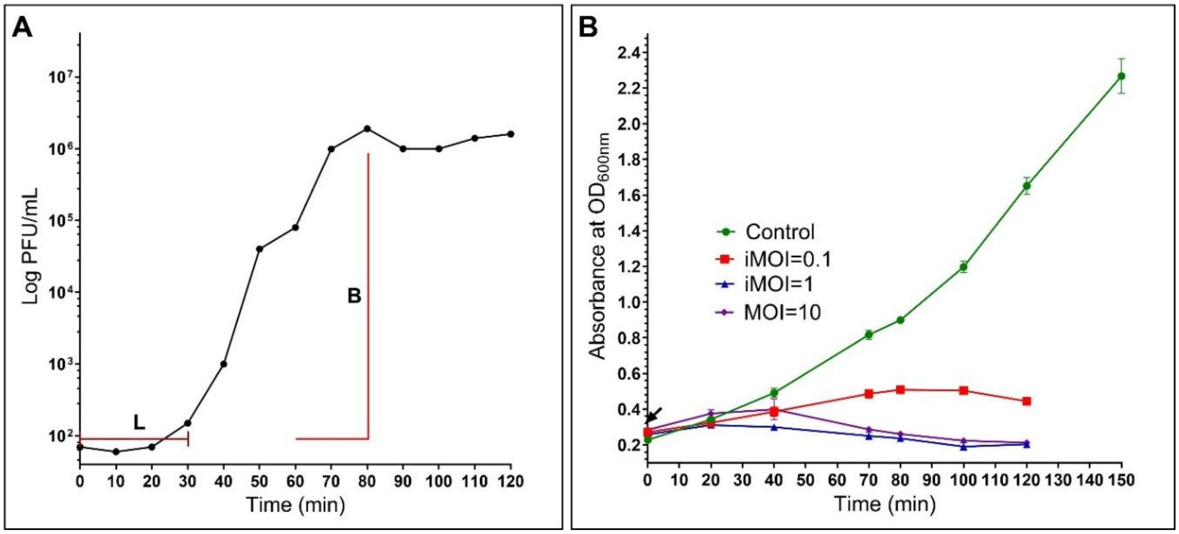


**Figure S5:** ***Pseudomonas*** **phage phiKZ growth kinetics. (A)** One-step growth curve of phiKZ using PAO1 as the host in LB medium at iMOI of 0.1. **(B)** The killing kinetics of phiKZ at iMOI of 0.1, 1.0 and 10 over a window of 2 hours in the shaker flask. The data represents the mean ± standard deviation of three independent experiments.

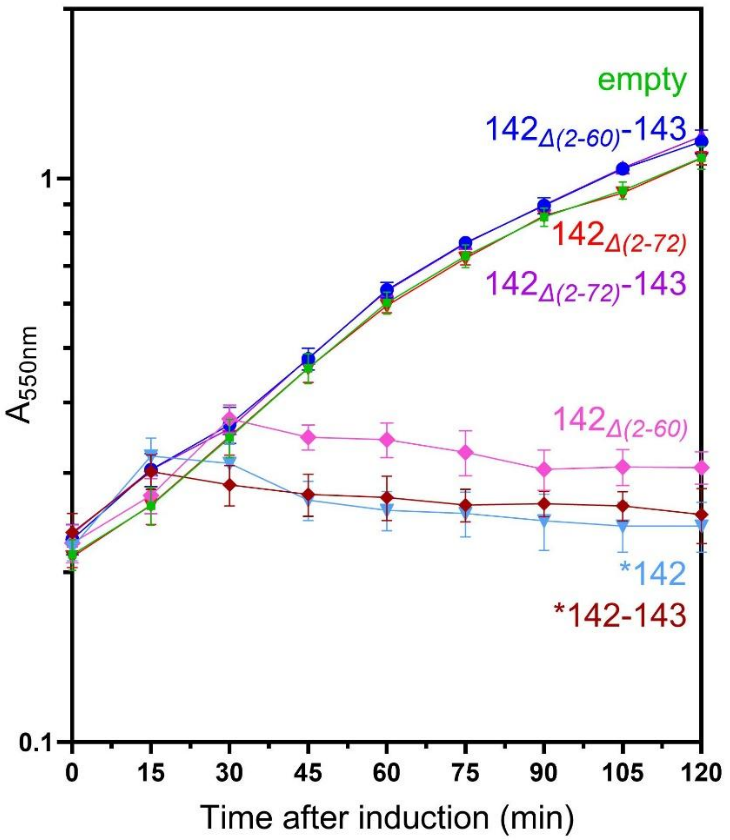


**Figure S6: Functional analysis of phiKZ holin mutant alleles.** In the presence of full length gp142, N-terminal binds with gp143 to control the lysis timing, that is, the binding fastens the holin activity. But the interaction of truncated N-terminal gp142 i.e., gp142$_{\Delta(2\text{-}60)}$ or gp142$_{\Delta(2\text{-}72)}$ has inhibited the holin-mediated lysis. This also confirms the interaction between N-terminal of gp142 with gp143. See figure 5 for mutant allele sequences. The mutant holin alleles with and without gp143 in pBAD vector were induced with arabinose in MG1655 cells in the following combinations: empty pBAD24 vector (blue), pBAD24-gp142 (maroon), pBAD24-gp142+143 (light blue), pBAD24- gp142$_{\Delta(2\text{-}60)}$ (pink), pBAD24- gp142$_{\Delta(2\text{-}29)}$+143 (blue), pBAD24- gp142$_{\Delta(2\text{-}72)}$ (red), and pBAD24- gp142$_{\Delta(2\text{-}72)}$+143 (purple). The data represents the mean ± standard deviation of three independent experiments.
