## Supplemental Data for "The Lysis Cassette of Jumbophage phiKZ"

**Supplementary material:**

**Supplementary data 1:** List of all the phiKZ genes with transmembrane domain (TMD). The results or output was generated using TMHMM webserver (https://services.healthtech.dtu.dk/services/TMHMM-2.0/).

**TMHMM result:**

### PHIKZ_p02 Length: 49
### PHIKZ_p02 Number of predicted TMHs: 0
### PHIKZ_p02 Exp number of AAs in TMHs: 8.99207
### PHIKZ_p02 Exp number, first 60 AAs: 8.99207
### PHIKZ_p02 Total prob of N-in: 0.21955
PHIKZ_p02 TMHMM2.0 outside 1 49

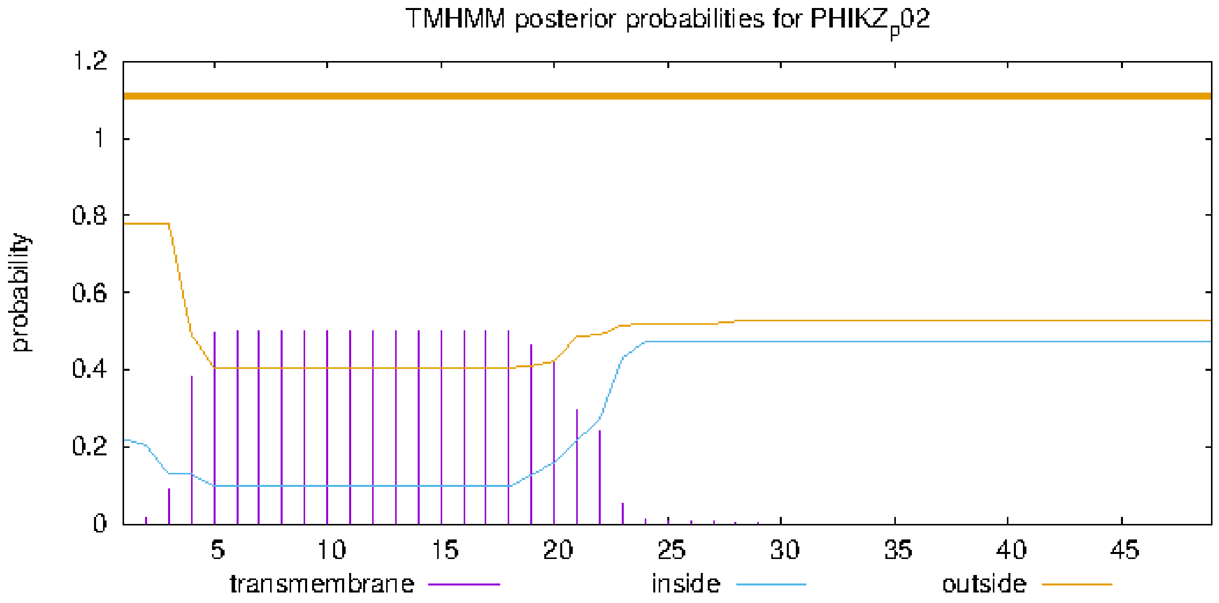


### plot in postscript, script for making the plot in gnuplot, data for plot

### PHIKZ012 Length: 132
### PHIKZ012 Number of predicted TMHs:  1
### PHIKZ012 Exp number of AAs in TMHs: 22.33116
### PHIKZ012 Exp number, first 60 AAs:  22.33075
# PHIKZ012 Total prob of N-in:        0.84338
### PHIKZ012 POSSIBLE N-term signal sequence

| | | | | |
|---|---|---|---|---|
| PHIKZ012 | TMHMM2.0 | inside | 1 | 6 |
| PHIKZ012 | TMHMM2.0 | TMhelix | 7 | 29 |
| PHIKZ012 | TMHMM2.0 | outside | 30 | 132 |

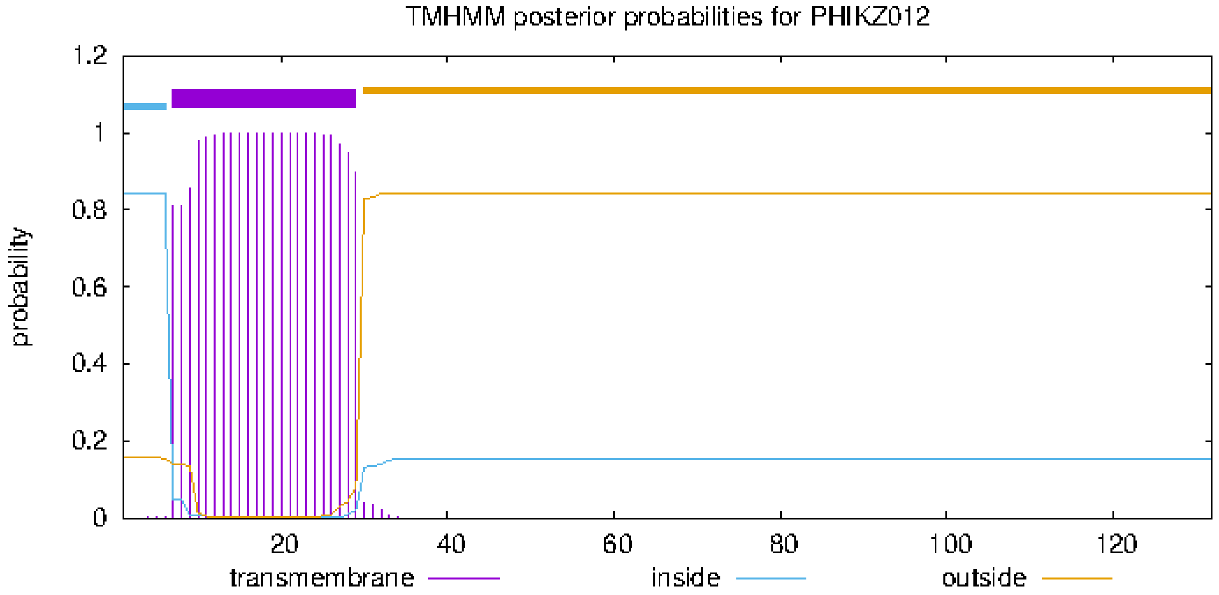


### plot in postscript, script for making the plot in gnuplot, data for plot

### PHIKZ019 Length: 69
### PHIKZ019 Number of predicted TMHs:  2
### PHIKZ019 Exp number of AAs in TMHs: 40.7431
### PHIKZ019 Exp number, first 60 AAs:  37.25197
# PHIKZ019 Total prob of N-in:        0.05539
### PHIKZ019 POSSIBLE N-term signal sequence

| | | | | |
|---|---|---|---|---|
| PHIKZ019 | TMHMM2.0 | outside | 1 | 3 |
| PHIKZ019 | TMHMM2.0 | TMhelix | 4 | 23 |
| PHIKZ019 | TMHMM2.0 | inside | 24 | 42 |
| PHIKZ019 | TMHMM2.0 | TMhelix | 43 | 65 |
| PHIKZ019 | TMHMM2.0 | outside | 66 | 69 |

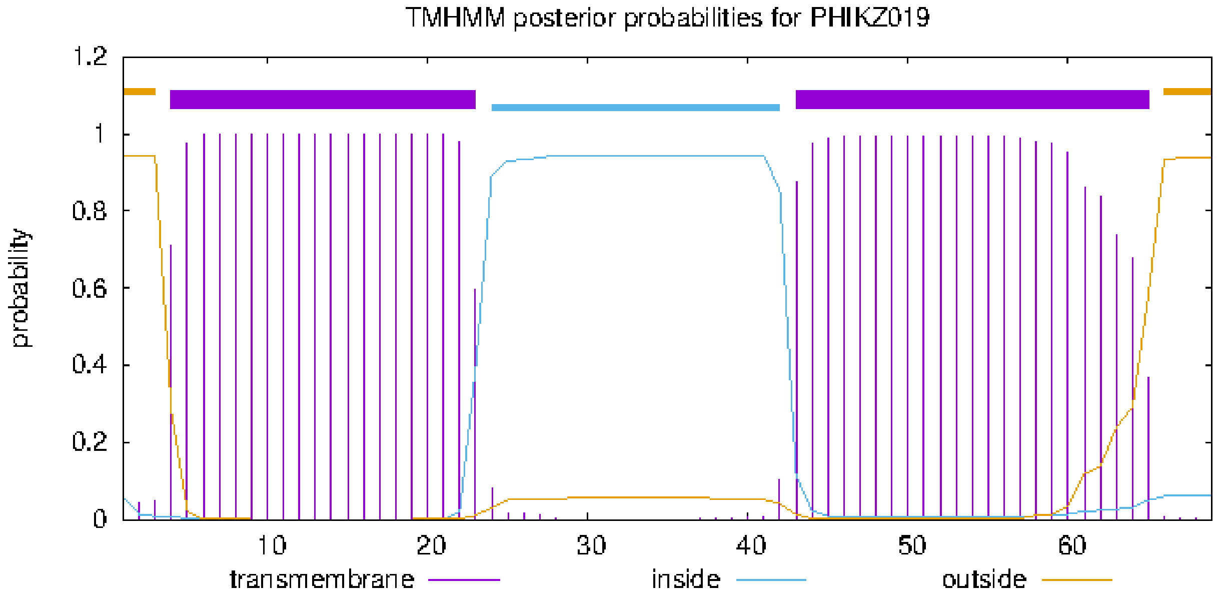


### plot in postscript, script for making the plot in gnuplot, data for plot

### PHIKZ023 Length: 138
### PHIKZ023 Number of predicted TMHs:  2
### PHIKZ023 Exp number of AAs in TMHs: 42.83242
### PHIKZ023 Exp number, first 60 AAs:  0.02319
# PHIKZ023 Total prob of N-in:        0.82463

| | | | | |
|---|---|---|---|---|
| PHIKZ023 | TMHMM2.0 | inside | 1 | 75 |
| PHIKZ023 | TMHMM2.0 | TMhelix | 76 | 98 |
| PHIKZ023 | TMHMM2.0 | outside | 99 | 107 |
| PHIKZ023 | TMHMM2.0 | TMhelix | 108 | 130 |
| PHIKZ023 | TMHMM2.0 | inside | 131 | 138 |

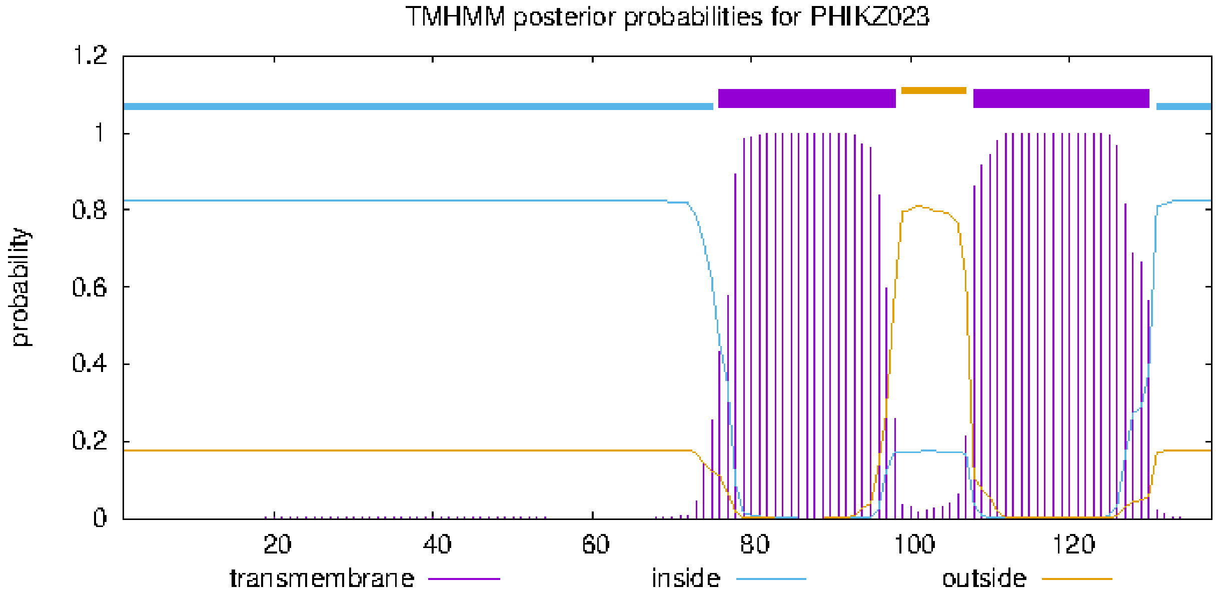


### plot in postscript, script for making the plot in gnuplot, data for plot

### PHIKZ_p08 Length: 412
### PHIKZ_p08 Number of predicted TMHs:  0
### PHIKZ_p08 Exp number of AAs in TMHs: 10.64769
### PHIKZ_p08 Exp number, first 60 AAs:  0.03325
# PHIKZ_p08 Total prob of N-in:        0.04045
PHIKZ_p08   TMHMM2.0   outside        1   412

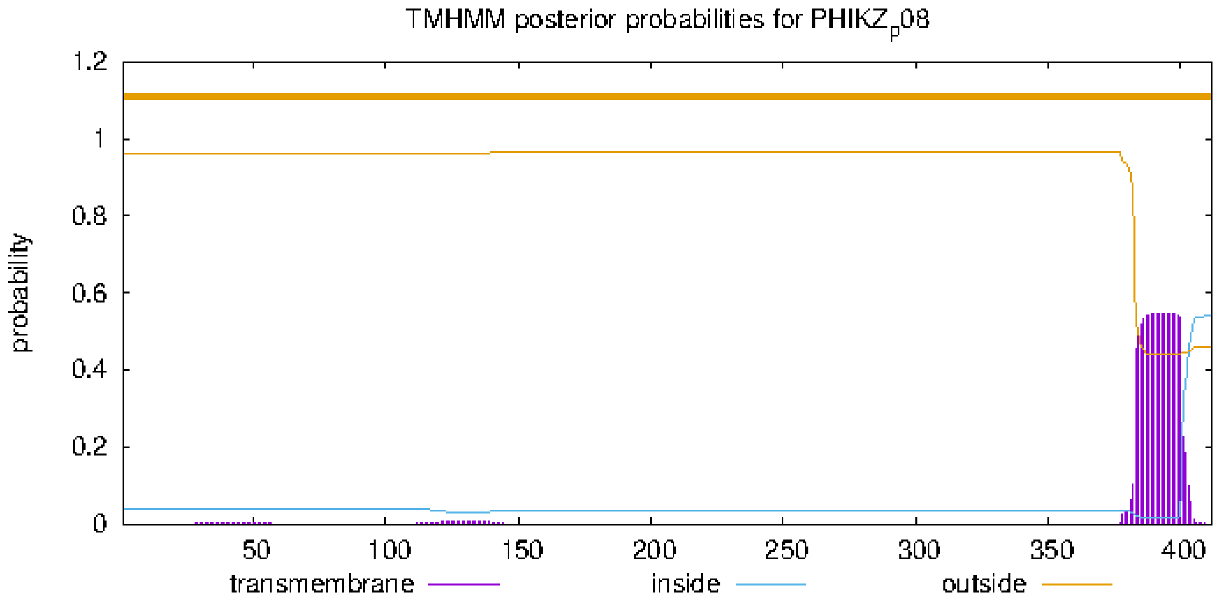


### plot in postscript, script for making the plot in gnuplot, data for plot

### PHIKZ076 Length: 85
### PHIKZ076 Number of predicted TMHs:  2
### PHIKZ076 Exp number of AAs in TMHs: 45.0976
### PHIKZ076 Exp number, first 60 AAs:  32.12435
# PHIKZ076 Total prob of N-in:        0.28597
### PHIKZ076 POSSIBLE N-term signal sequence

| | | | | |
|---|---|---|---|---|
| PHIKZ076 | TMHMM2.0 | outside | 1 | 9 |
| PHIKZ076 | TMHMM2.0 | TMhelix | 10 | 32 |
| PHIKZ076 | TMHMM2.0 | inside | 33 | 51 |
| PHIKZ076 | TMHMM2.0 | TMhelix | 52 | 74 |
| PHIKZ076 | TMHMM2.0 | outside | 75 | 85 |

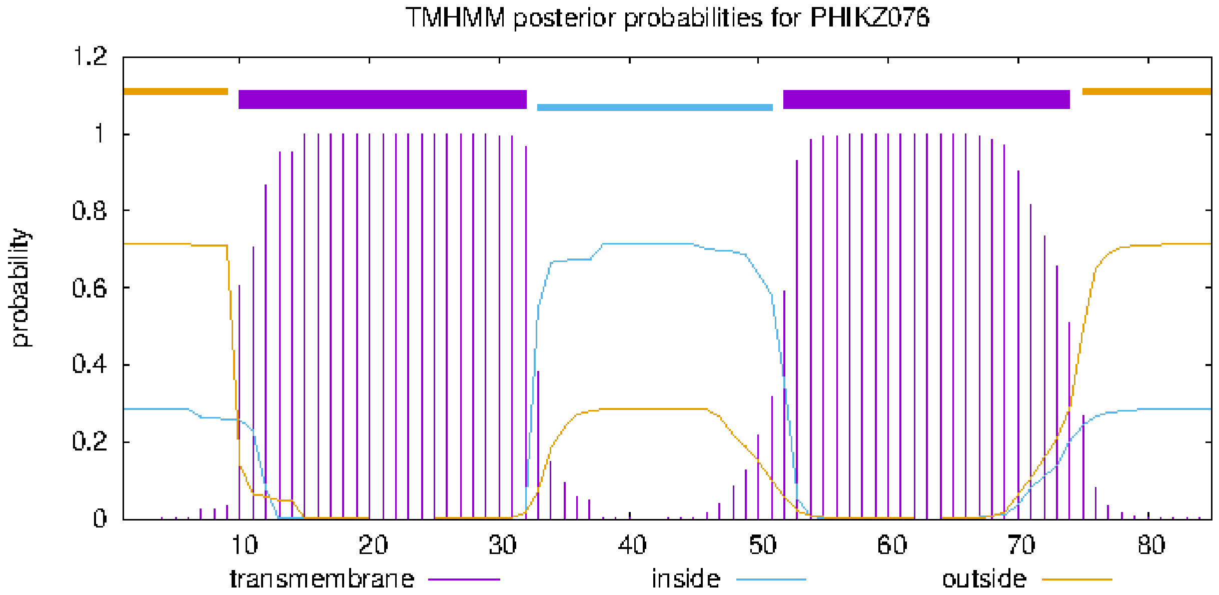


### plot in postscript, script for making the plot in gnuplot, data for plot

### PHIKZ_p23 Length: 63
### PHIKZ_p23 Number of predicted TMHs:  2
### PHIKZ_p23 Exp number of AAs in TMHs: 45.355
### PHIKZ_p23 Exp number, first 60 AAs:  45.17631
# PHIKZ_p23 Total prob of N-in:        0.63621
### PHIKZ_p23 POSSIBLE N-term signal sequence

| | | | | |
|---|---|---|---|---|
| PHIKZ_p23 | TMHMM2.0 | inside | 1 | 4 |
| PHIKZ_p23 | TMHMM2.0 | TMhelix | 5 | 27 |
| PHIKZ_p23 | TMHMM2.0 | outside | 28 | 36 |
| PHIKZ_p23 | TMHMM2.0 | TMhelix | 37 | 59 |
| PHIKZ_p23 | TMHMM2.0 | inside | 60 | 63 |

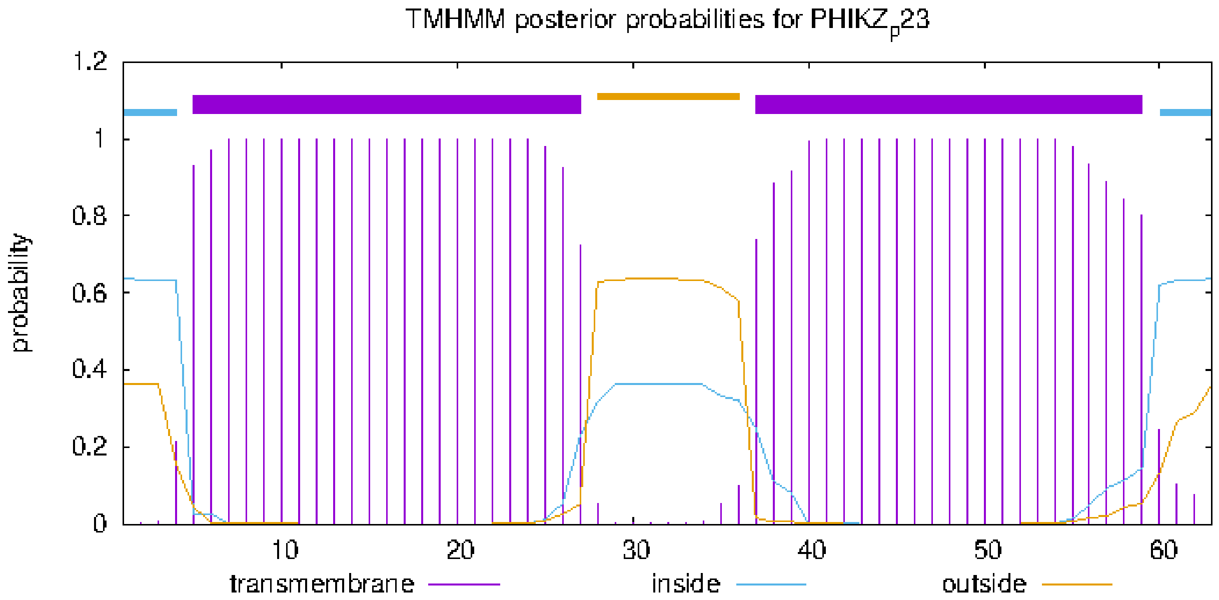


### plot in postscript, script for making the plot in gnuplot, data for plot

### PHIKZ_p24 Length: 64
### PHIKZ_p24 Number of predicted TMHs: 2
### PHIKZ_p24 Exp number of AAs in TMHs: 39.56245
### PHIKZ_p24 Exp number, first 60 AAs: 36.90403
### PHIKZ_p24 Total prob of N-in: 0.30611
### PHIKZ_p24 POSSIBLE N-term signal sequence

| | | | | |
|---|---|---|---|---|
| PHIKZ_p24 | TMHMM2.0 | outside | 1 | 3 |
| PHIKZ_p24 | TMHMM2.0 | TMhelix | 4 | 26 |
| PHIKZ_p24 | TMHMM2.0 | inside | 27 | 45 |
| PHIKZ_p24 | TMHMM2.0 | TMhelix | 46 | 63 |
| PHIKZ_p24 | TMHMM2.0 | outside | 64 | 64 |

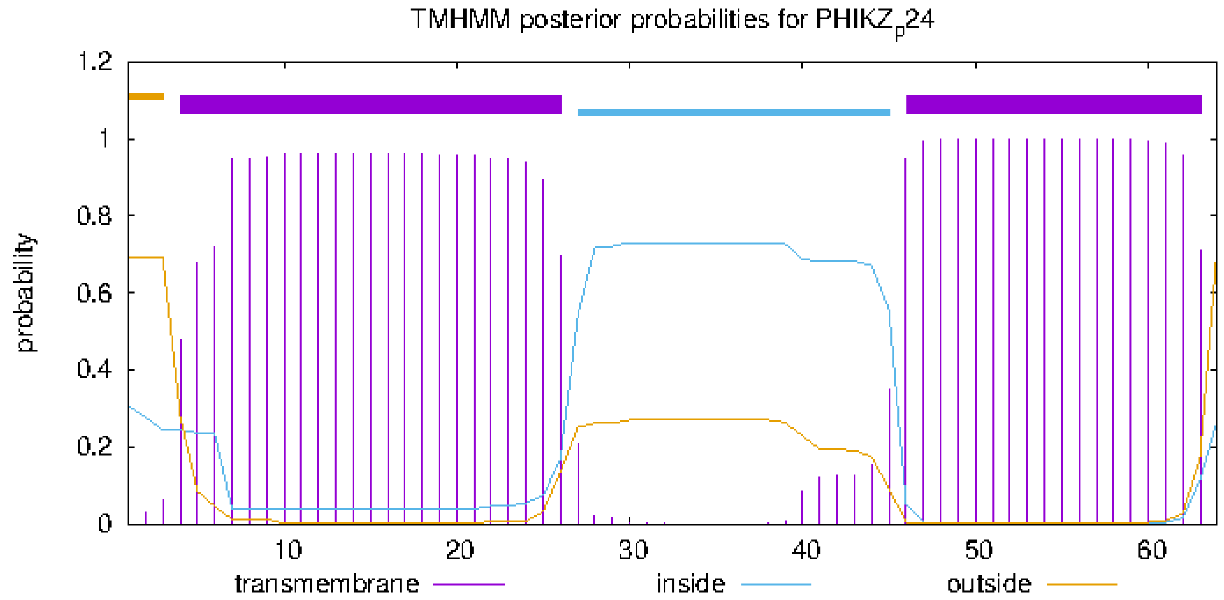


### plot in postscript, script for making the plot in gnuplot, data for plot

### PHIKZ_p32 Length: 41
### PHIKZ_p32 Number of predicted TMHs:  1
### PHIKZ_p32 Exp number of AAs in TMHs: 18.87973
### PHIKZ_p32 Exp number, first 60 AAs:  18.87973
# PHIKZ_p32 Total prob of N-in:        0.77938
### PHIKZ_p32 POSSIBLE N-term signal sequence

| | | | | |
|---|---|---|---|---|
| PHIKZ_p32 | TMHMM2.0 | inside | 1 | 20 |
| PHIKZ_p32 | TMHMM2.0 | TMhelix | 21 | 38 |
| PHIKZ_p32 | TMHMM2.0 | outside | 39 | 41 |

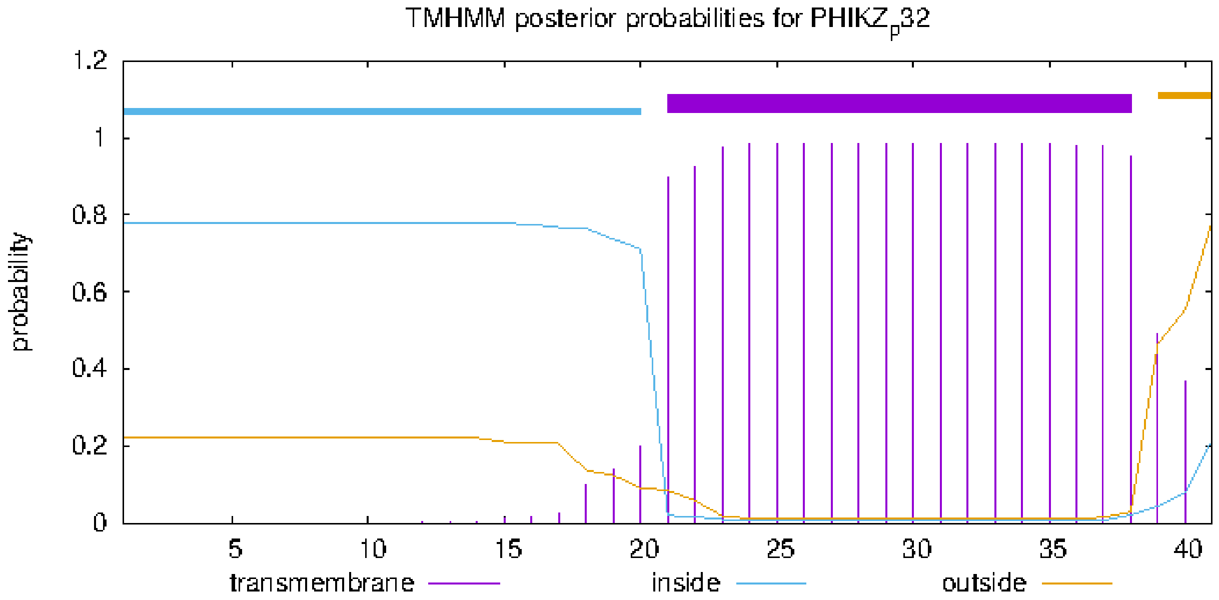


### plot in postscript, script for making the plot in gnuplot, data for plot

### PHIKZ103 Length: 130
### PHIKZ103 Number of predicted TMHs:  1
### PHIKZ103 Exp number of AAs in TMHs: 20.72715
### PHIKZ103 Exp number, first 60 AAs:  20.72379
# PHIKZ103 Total prob of N-in:        0.55831
### PHIKZ103 POSSIBLE N-term signal sequence

| | | | | |
|---|---|---|---|---|
| PHIKZ103 | TMHMM2.0 | outside | 1 | 3 |
| PHIKZ103 | TMHMM2.0 | TMhelix | 4 | 26 |
| PHIKZ103 | TMHMM2.0 | inside | 27 | 130 |

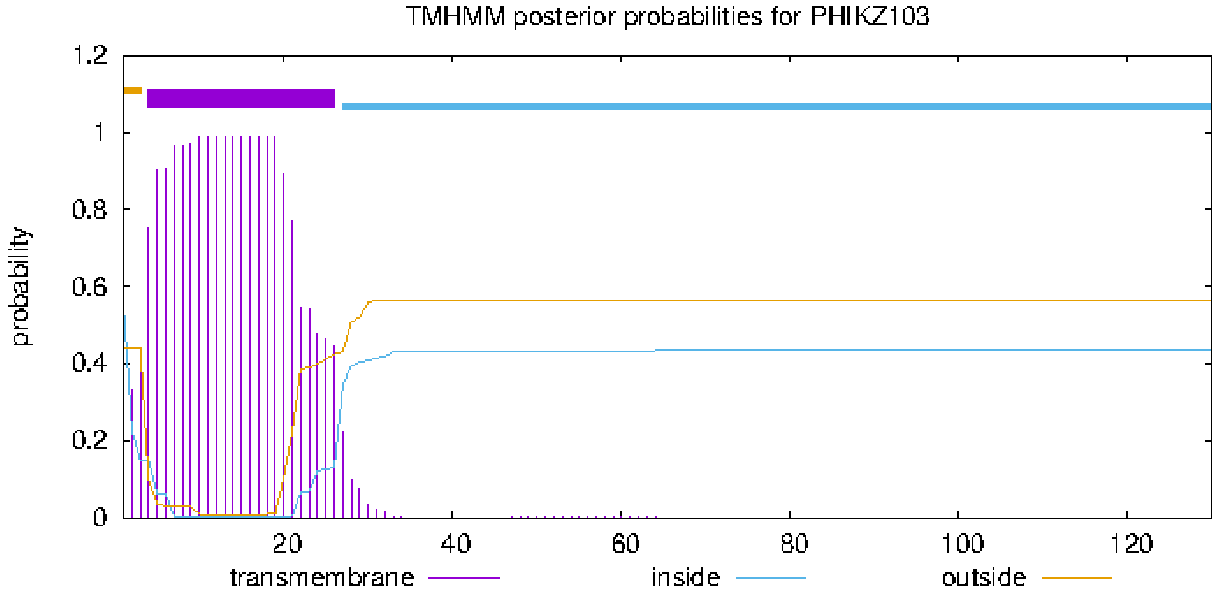


### plot in postscript, script for making the plot in gnuplot, data for plot

### PHIKZ106 Length: 112
### PHIKZ106 Number of predicted TMHs:  1
### PHIKZ106 Exp number of AAs in TMHs: 20.42495
### PHIKZ106 Exp number, first 60 AAs:  20.4225
# PHIKZ106 Total prob of N-in:        0.08861
### PHIKZ106 POSSIBLE N-term signal sequence
PHIKZ106     TMHMM2.0   outside      1     4
PHIKZ106     TMHMM2.0   TMhelix      5    27
PHIKZ106     TMHMM2.0   inside      28   112

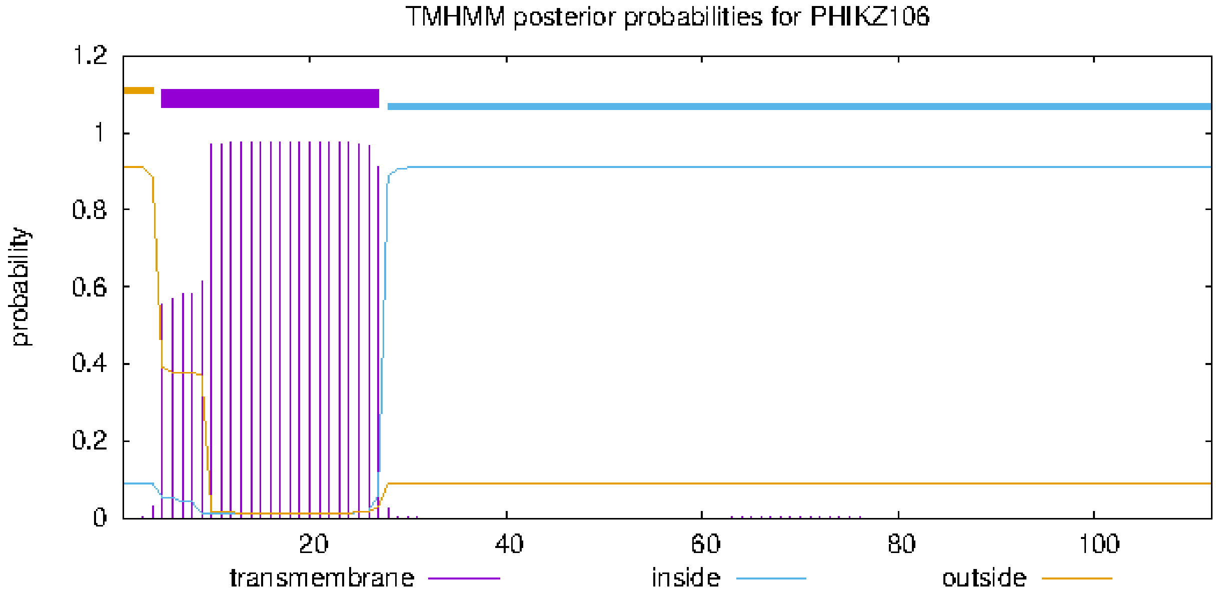


### plot in postscript, script for making the plot in gnuplot, data for plot

### PHIKZ_p39 Length: 69
### PHIKZ_p39 Number of predicted TMHs: 1
### PHIKZ_p39 Exp number of AAs in TMHs: 23.34415
### PHIKZ_p39 Exp number, first 60 AAs: 22.69827
### PHIKZ_p39 Total prob of N-in: 0.72939
### PHIKZ_p39 POSSIBLE N-term signal sequence

| | | | | |
|---|---|---|---|---|
| PHIKZ_p39 | TMHMM2.0 | inside | 1 | 37 |
| PHIKZ_p39 | TMHMM2.0 | TMhelix | 38 | 60 |
| PHIKZ_p39 | TMHMM2.0 | outside | 61 | 69 |

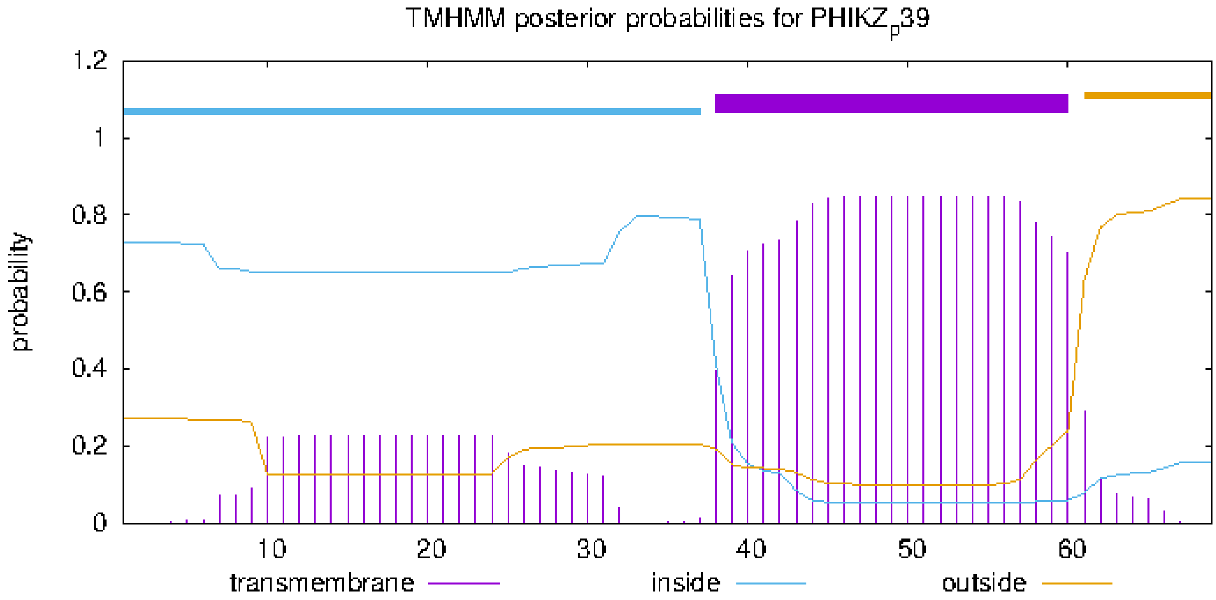


### plot in postscript, script for making the plot in gnuplot, data for plot

### PHIKZ_p41 Length: 78
### PHIKZ_p41 Number of predicted TMHs:  1
### PHIKZ_p41 Exp number of AAs in TMHs: 22.71605
### PHIKZ_p41 Exp number, first 60 AAs:  22.69346
# PHIKZ_p41 Total prob of N-in:        0.46647
### PHIKZ_p41 POSSIBLE N-term signal sequence

PHIKZ_p41   TMHMM2.0   outside   1   5
PHIKZ_p41   TMHMM2.0   TMhelix   6   28
PHIKZ_p41   TMHMM2.0   inside   29   78

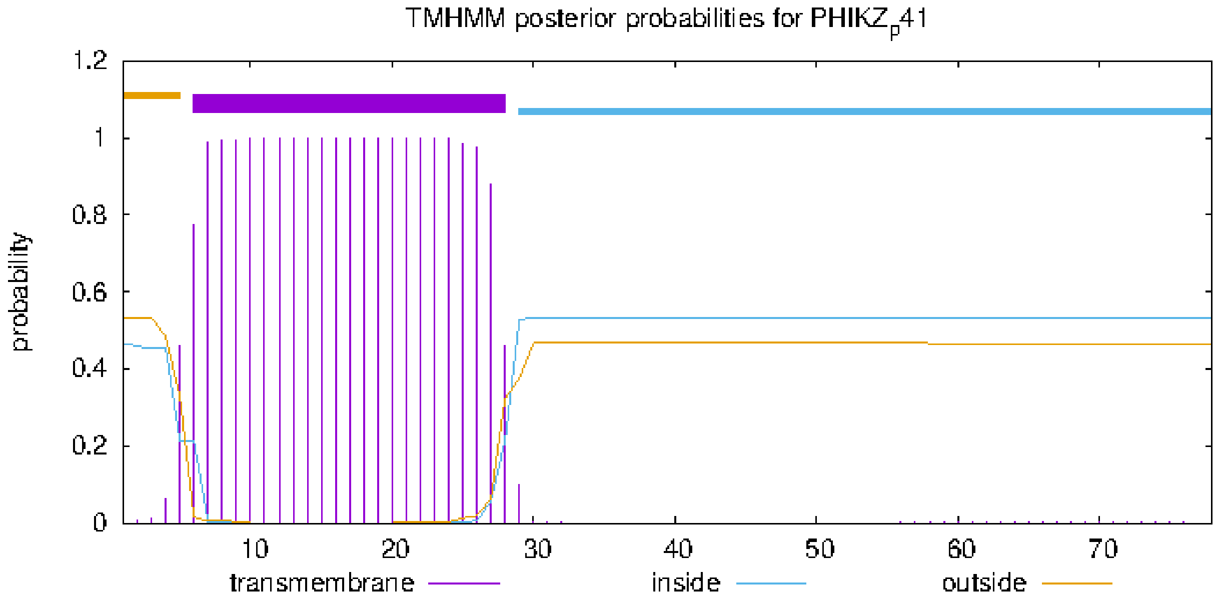


### plot in postscript, script for making the plot in gnuplot, data for plot

### PHIKZ141 Length: 168
### PHIKZ141 Number of predicted TMHs: 1
### PHIKZ141 Exp number of AAs in TMHs: 22.47823
### PHIKZ141 Exp number, first 60 AAs: 22.47823
### PHIKZ141 Total prob of N-in: 0.50438
### PHIKZ141 POSSIBLE N-term signal sequence

| | | | | |
|---|---|---|---|---|
| PHIKZ141 | TMHMM2.0 | inside | 1 | 12 |
| PHIKZ141 | TMHMM2.0 | TMhelix | 13 | 35 |
| PHIKZ141 | TMHMM2.0 | outside | 36 | 168 |

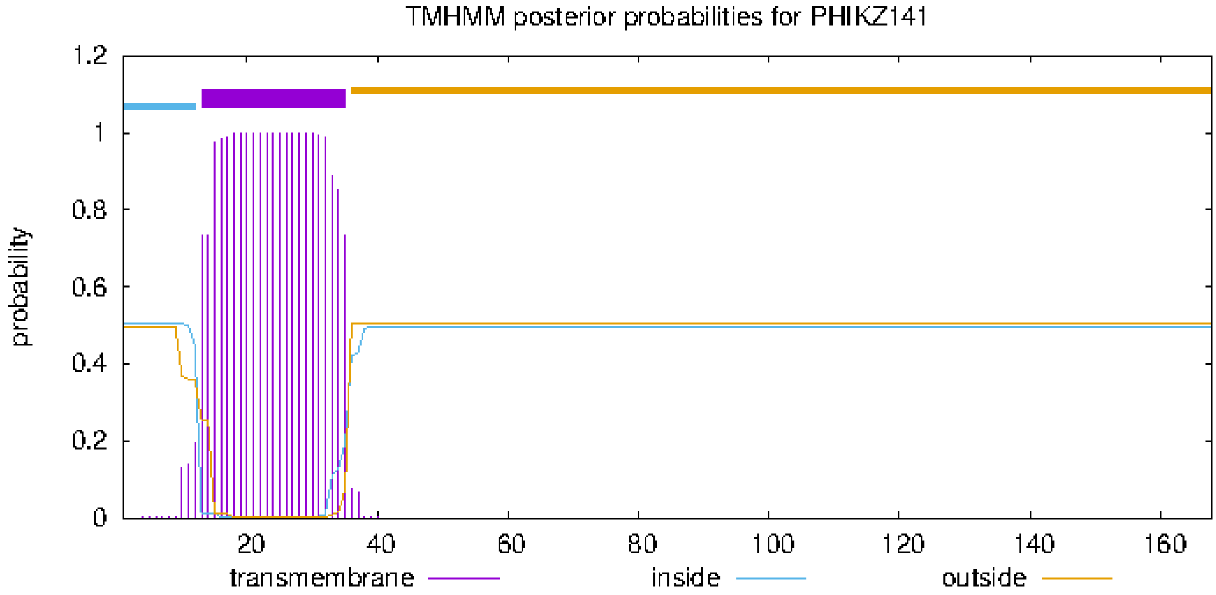


### plot in postscript, script for making the plot in gnuplot, data for plot

### PHIKZ142 Length: 131
### PHIKZ142 Number of predicted TMHs:  1
### PHIKZ142 Exp number of AAs in TMHs: 22.10504
### PHIKZ142 Exp number, first 60 AAs:  0.01192
# PHIKZ142 Total prob of N-in:        0.94186

| | | | | |
|---|---|---|---|---|
| PHIKZ142 | TMHMM2.0 | inside | 1 | 80 |
| PHIKZ142 | TMHMM2.0 | TMhelix | 81 | 103 |
| PHIKZ142 | TMHMM2.0 | outside | 104 | 131 |

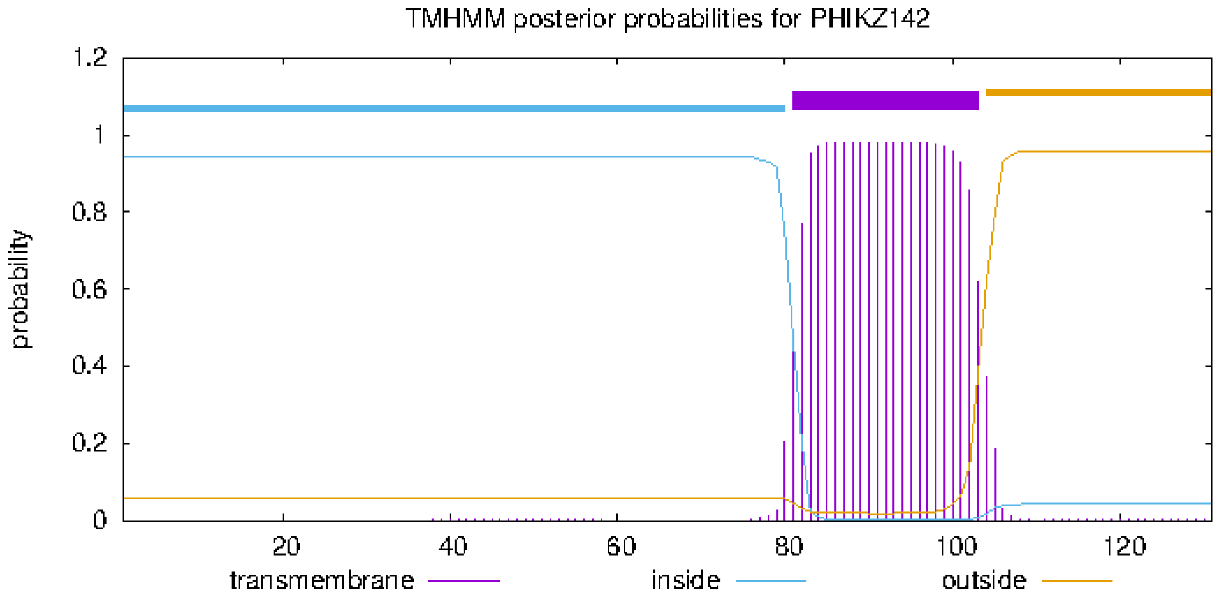


### plot in postscript, script for making the plot in gnuplot, data for plot

### PHIKZ_p47 Length: 96
### PHIKZ_p47 Number of predicted TMHs:  1
### PHIKZ_p47 Exp number of AAs in TMHs: 21.57482
### PHIKZ_p47 Exp number, first 60 AAs:  21.56691
# PHIKZ_p47 Total prob of N-in:        0.15356
### PHIKZ_p47 POSSIBLE N-term signal sequence

| | | | | |
|---|---|---|---|---|
| PHIKZ_p47 | TMHMM2.0 | outside | 1 | 35 |
| PHIKZ_p47 | TMHMM2.0 | TMhelix | 36 | 55 |
| PHIKZ_p47 | TMHMM2.0 | inside | 56 | 96 |

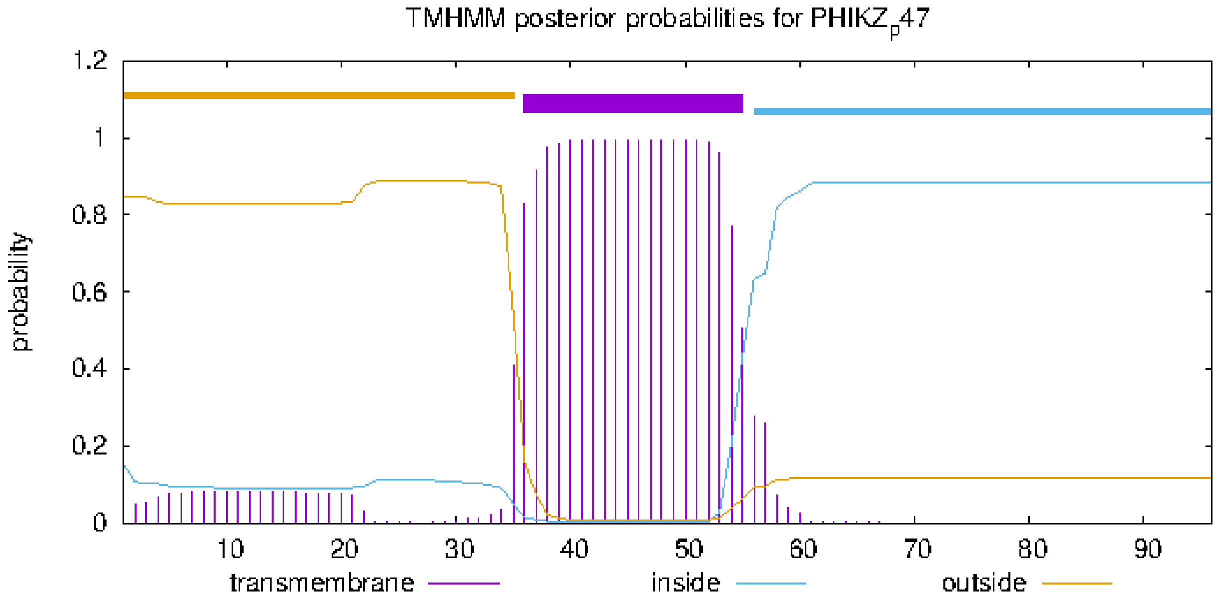


### plot in postscript, script for making the plot in gnuplot, data for plot

### PHIKZ_p53 Length: 68
### PHIKZ_p53 Number of predicted TMHs:  1
### PHIKZ_p53 Exp number of AAs in TMHs: 16.63289
### PHIKZ_p53 Exp number, first 60 AAs:  16.63289
# PHIKZ_p53 Total prob of N-in:        0.30684
### PHIKZ_p53 POSSIBLE N-term signal sequence

PHIKZ_p53	TMHMM2.0	outside	1	5
PHIKZ_p53	TMHMM2.0	TMhelix	6	28
PHIKZ_p53	TMHMM2.0	inside	29	68

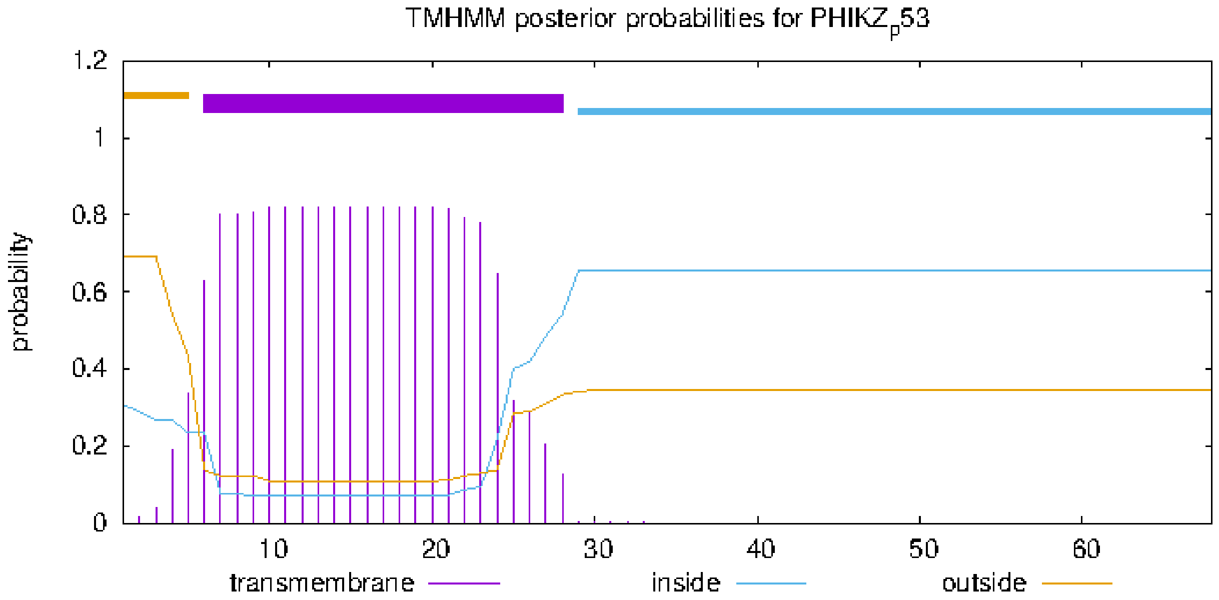


### plot in postscript, script for making the plot in gnuplot, data for plot

### PHIKZ167 Length: 142
### PHIKZ167 Number of predicted TMHs:  1
### PHIKZ167 Exp number of AAs in TMHs: 18.82654
### PHIKZ167 Exp number, first 60 AAs:  0.00773
# PHIKZ167 Total prob of N-in:        0.03667
PHIKZ167     TMHMM2.0   outside          1   115
PHIKZ167     TMHMM2.0   TMhelix        116   135
PHIKZ167     TMHMM2.0   inside     136   142

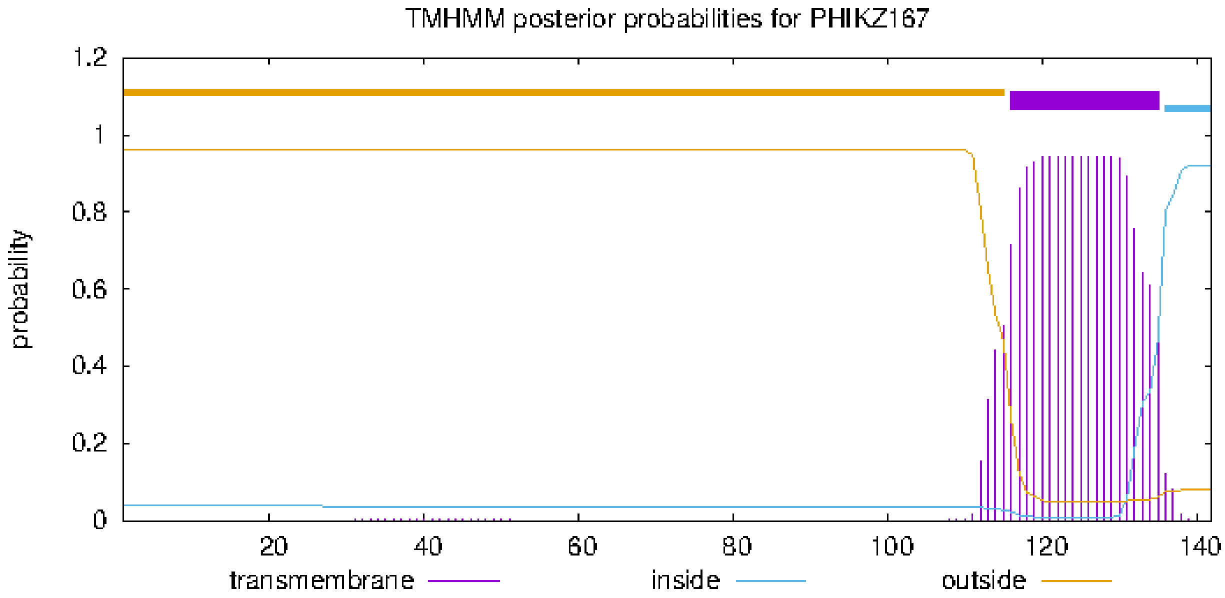


### plot in postscript, script for making the plot in gnuplot, data for plot

### PHIKZ197 Length: 120
### PHIKZ197 Number of predicted TMHs:  3
### PHIKZ197 Exp number of AAs in TMHs: 64.17528
### PHIKZ197 Exp number, first 60 AAs:  24.75631
# PHIKZ197 Total prob of N-in:        0.97443
### PHIKZ197 POSSIBLE N-term signal sequence

| | | | | |
|---|---|---|---|---|
| PHIKZ197 | TMHMM2.0 | inside | 1 | 2 |
| PHIKZ197 | TMHMM2.0 | TMhelix | 3 | 22 |
| PHIKZ197 | TMHMM2.0 | outside | 23 | 54 |
| PHIKZ197 | TMHMM2.0 | TMhelix | 55 | 77 |
| PHIKZ197 | TMHMM2.0 | inside | 78 | 96 |
| PHIKZ197 | TMHMM2.0 | TMhelix | 97 | 119 |
| PHIKZ197 | TMHMM2.0 | outside | 120 | 120 |

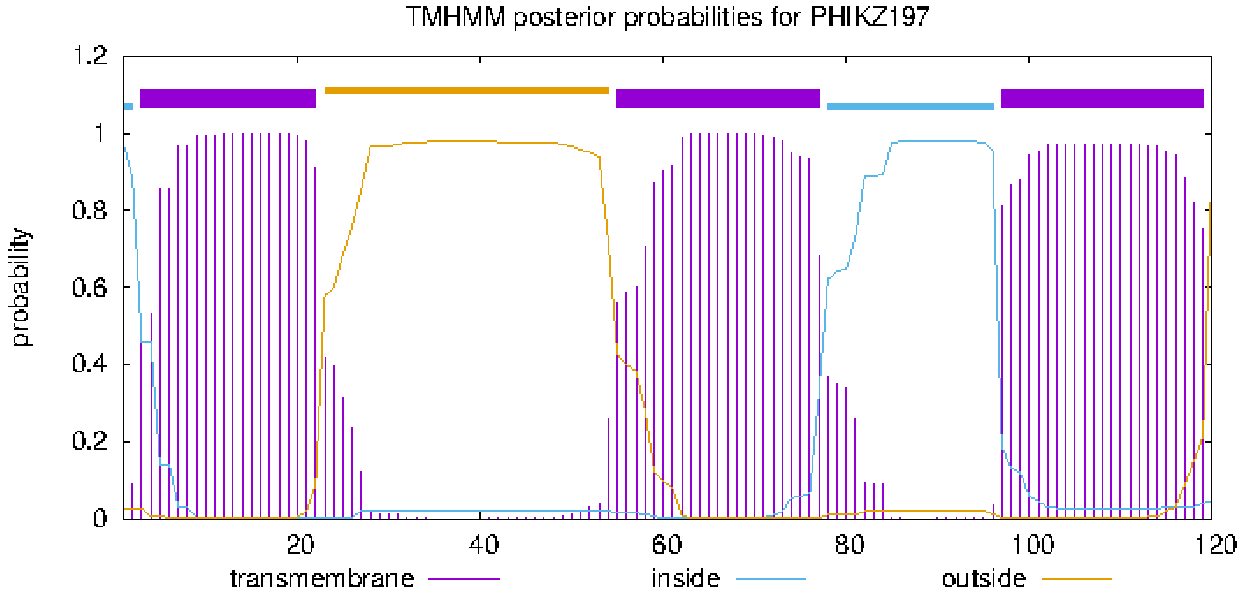


### plot in postscript, script for making the plot in gnuplot, data for plot

### PHIKZ226 Length: 109
### PHIKZ226 Number of predicted TMHs:  1
### PHIKZ226 Exp number of AAs in TMHs: 19.36797
### PHIKZ226 Exp number, first 60 AAs:  19.35726
# PHIKZ226 Total prob of N-in:        0.21989
### PHIKZ226 POSSIBLE N-term signal sequence

| | | | | |
|---|---|---|---|---|
| PHIKZ226 | TMHMM2.0 | outside | 1 | 3 |
| PHIKZ226 | TMHMM2.0 | TMhelix | 4 | 26 |
| PHIKZ226 | TMHMM2.0 | inside | 27 | 109 |

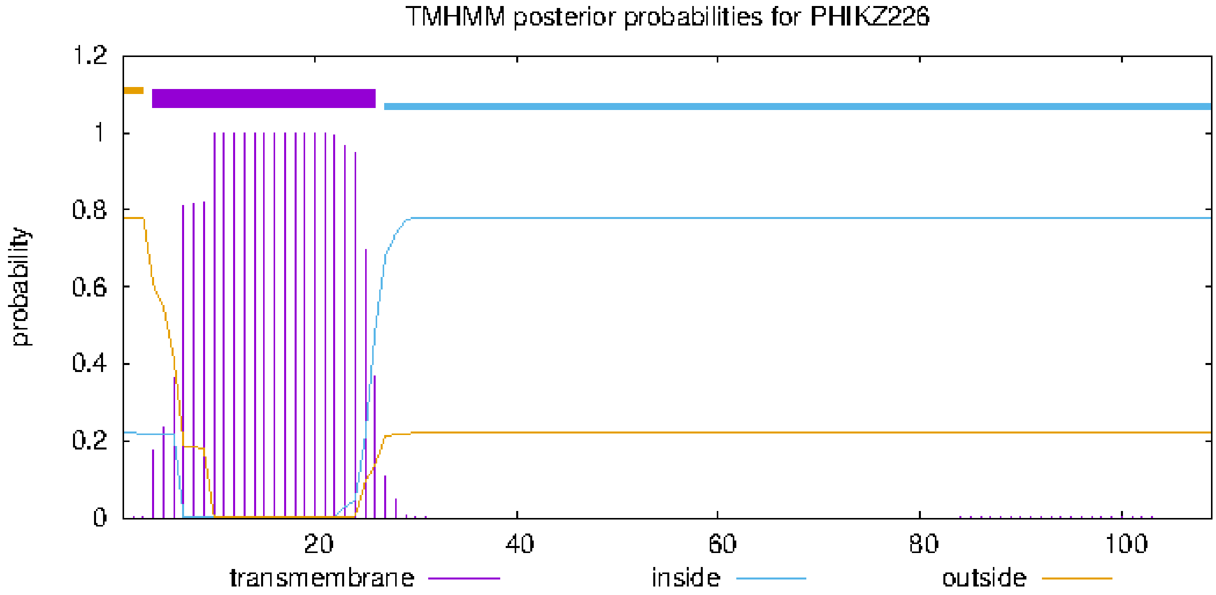


### plot in postscript, script for making the plot in gnuplot, data for plot

### PHIKZ_p65 Length: 98
### PHIKZ_p65 Number of predicted TMHs:  1
### PHIKZ_p65 Exp number of AAs in TMHs: 21.20508
### PHIKZ_p65 Exp number, first 60 AAs:  21.00794
# PHIKZ_p65 Total prob of N-in:        0.09543
### PHIKZ_p65 POSSIBLE N-term signal sequence
PHIKZ_p65   TMHMM2.0   outside   1   3
PHIKZ_p65   TMHMM2.0   TMhelix   4   23
PHIKZ_p65   TMHMM2.0   inside   24   98

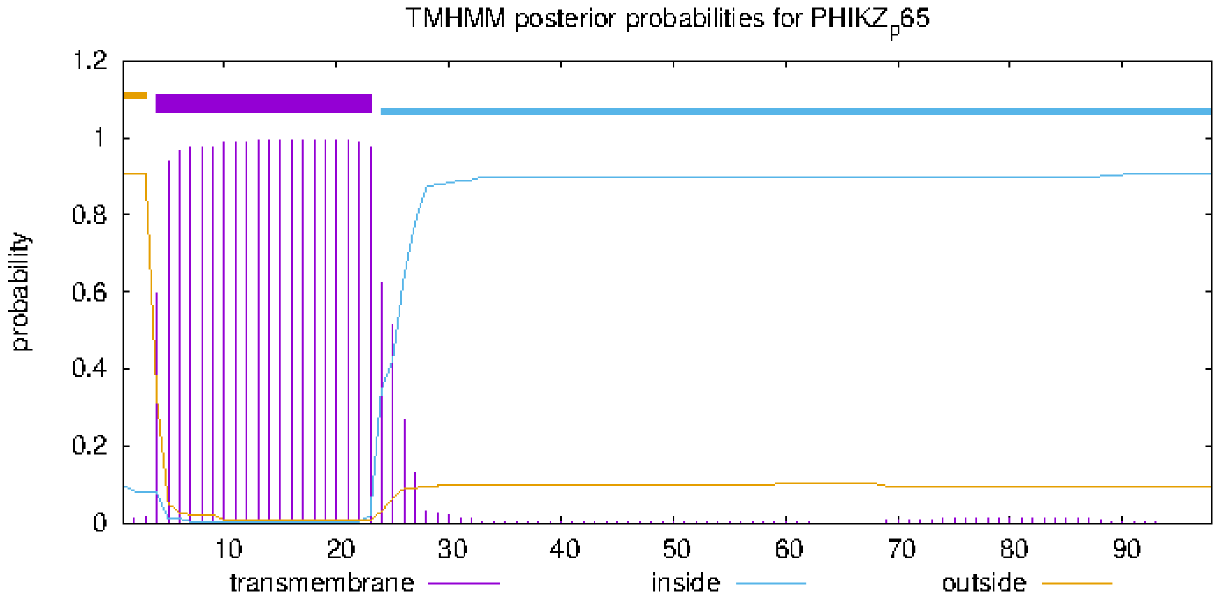


### plot in postscript, script for making the plot in gnuplot, data for plot

### PHIKZ_p66 Length: 96
### PHIKZ_p66 Number of predicted TMHs:  1
### PHIKZ_p66 Exp number of AAs in TMHs: 22.93172
### PHIKZ_p66 Exp number, first 60 AAs:  22.92657
# PHIKZ_p66 Total prob of N-in:        0.00541
### PHIKZ_p66 POSSIBLE N-term signal sequence

| | | | | |
|---|---|---|---|---|
| PHIKZ_p66 | TMHMM2.0 | outside | 1 | 3 |
| PHIKZ_p66 | TMHMM2.0 | TMhelix | 4 | 26 |
| PHIKZ_p66 | TMHMM2.0 | inside | 27 | 96 |

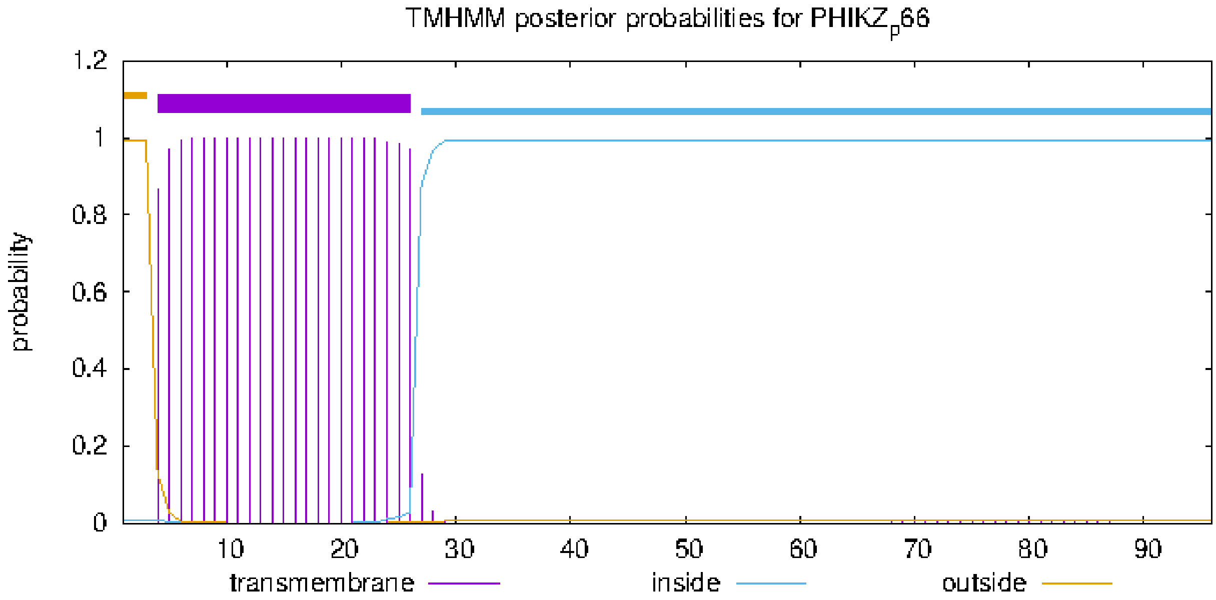


### plot in postscript, script for making the plot in gnuplot, data for plot

### PHIKZ_p67 Length: 57
### PHIKZ_p67 Number of predicted TMHs:  1
### PHIKZ_p67 Exp number of AAs in TMHs: 21.40061
### PHIKZ_p67 Exp number, first 60 AAs:  21.40061
# PHIKZ_p67 Total prob of N-in:        0.48176
### PHIKZ_p67 POSSIBLE N-term signal sequence
PHIKZ_p67   TMHMM2.0   outside      1     3
PHIKZ_p67   TMHMM2.0   TMhelix      4    26
PHIKZ_p67   TMHMM2.0   inside      27    57

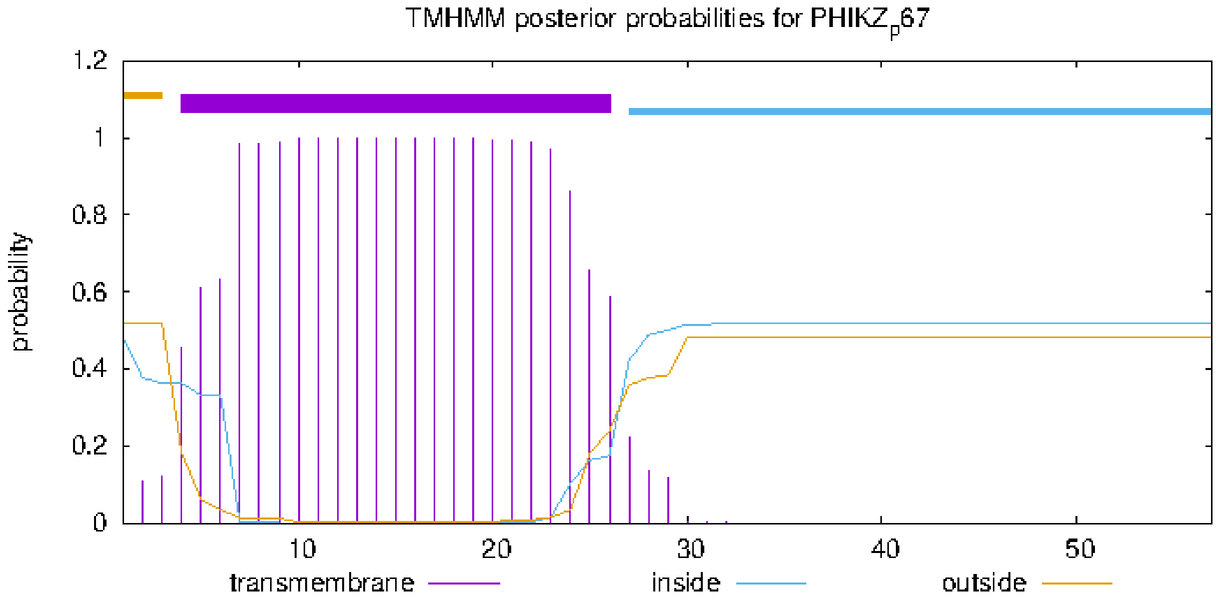


### plot in postscript, script for making the plot in gnuplot, data for plot

### PHIKZ241 Length: 109
### PHIKZ241 Number of predicted TMHs:  1
### PHIKZ241 Exp number of AAs in TMHs: 18.12134
### PHIKZ241 Exp number, first 60 AAs:  18.11825
# PHIKZ241 Total prob of N-in:        0.13949
### PHIKZ241 POSSIBLE N-term signal sequence

| | | | | |
|---|---|---|---|---|
| PHIKZ241 | TMHMM2.0 | outside | 1 | 3 |
| PHIKZ241 | TMHMM2.0 | TMhelix | 4 | 23 |
| PHIKZ241 | TMHMM2.0 | inside | 24 | 109 |

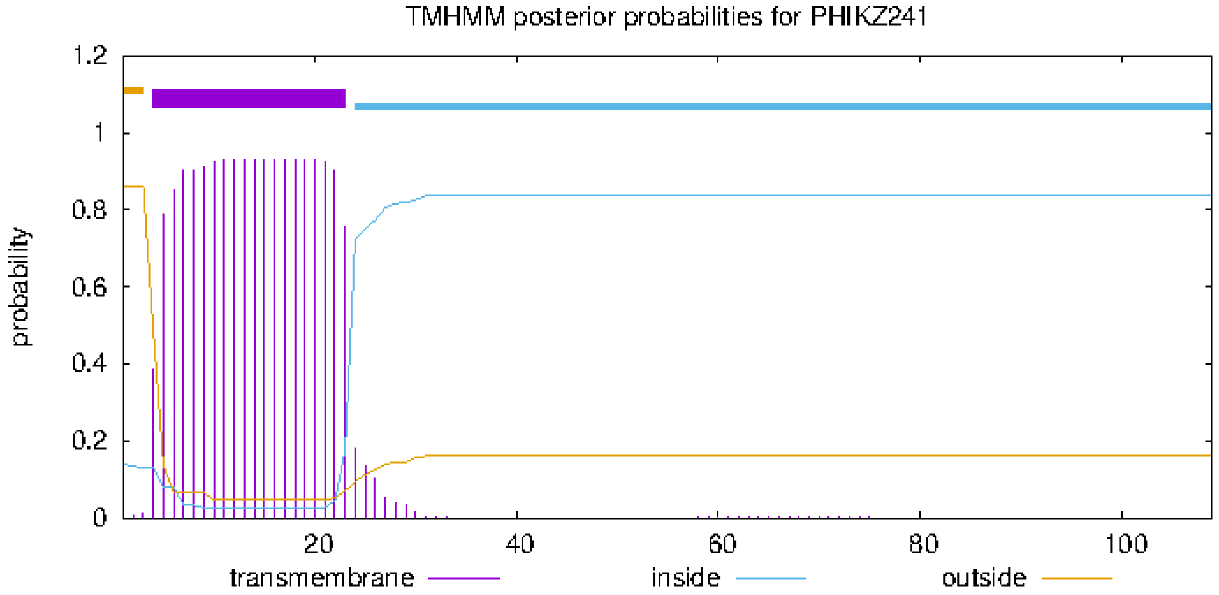


### plot in postscript, script for making the plot in gnuplot, data for plot

### PHIKZ249 Length: 264
### PHIKZ249 Number of predicted TMHs:  1
### PHIKZ249 Exp number of AAs in TMHs: 35.52726
### PHIKZ249 Exp number, first 60 AAs:  22.36436
# PHIKZ249 Total prob of N-in:        0.29877
### PHIKZ249 POSSIBLE N-term signal sequence

| | | | | |
|---|---|---|---|---|
| PHIKZ249 | TMHMM2.0 | inside | 1 | 4 |
| PHIKZ249 | TMHMM2.0 | TMhelix | 5 | 27 |
| PHIKZ249 | TMHMM2.0 | outside | 28 | 264 |

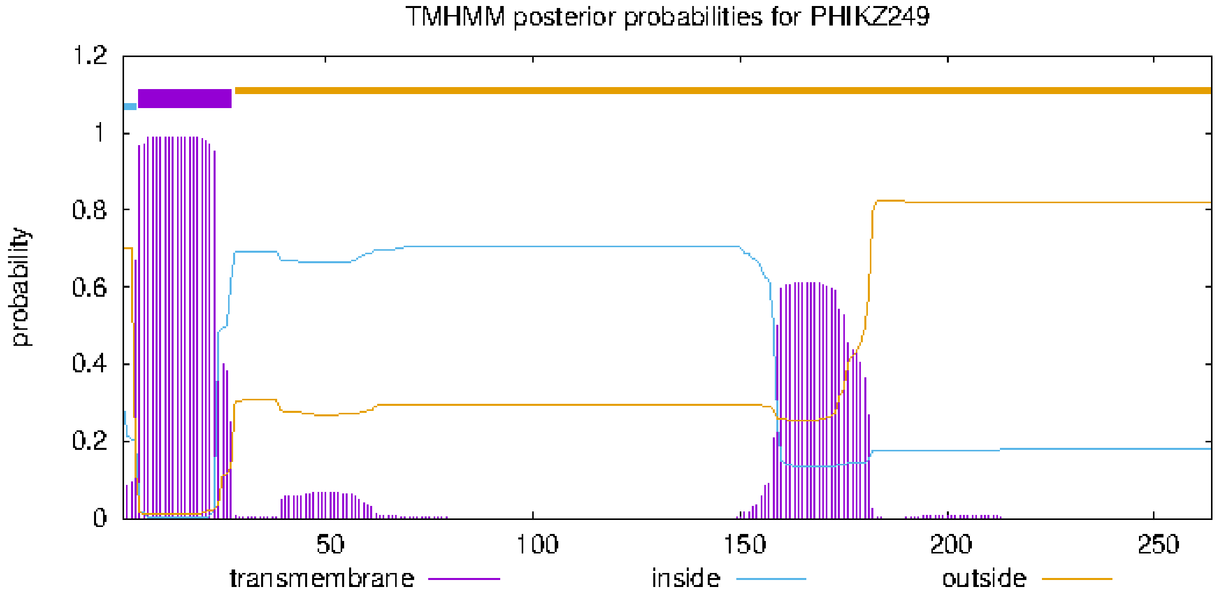


### plot in postscript, script for making the plot in gnuplot, data for plot

### PHIKZ251 Length: 151
### PHIKZ251 Number of predicted TMHs:  1
### PHIKZ251 Exp number of AAs in TMHs: 21.95435
### PHIKZ251 Exp number, first 60 AAs:  20.7852
# PHIKZ251 Total prob of N-in:        0.35000
### PHIKZ251 POSSIBLE N-term signal sequence

| | | | | |
|---|---|---|---|---|
| PHIKZ251 | TMHMM2.0 | outside | 1 | 3 |
| PHIKZ251 | TMHMM2.0 | TMhelix | 4 | 23 |
| PHIKZ251 | TMHMM2.0 | inside | 24 | 151 |

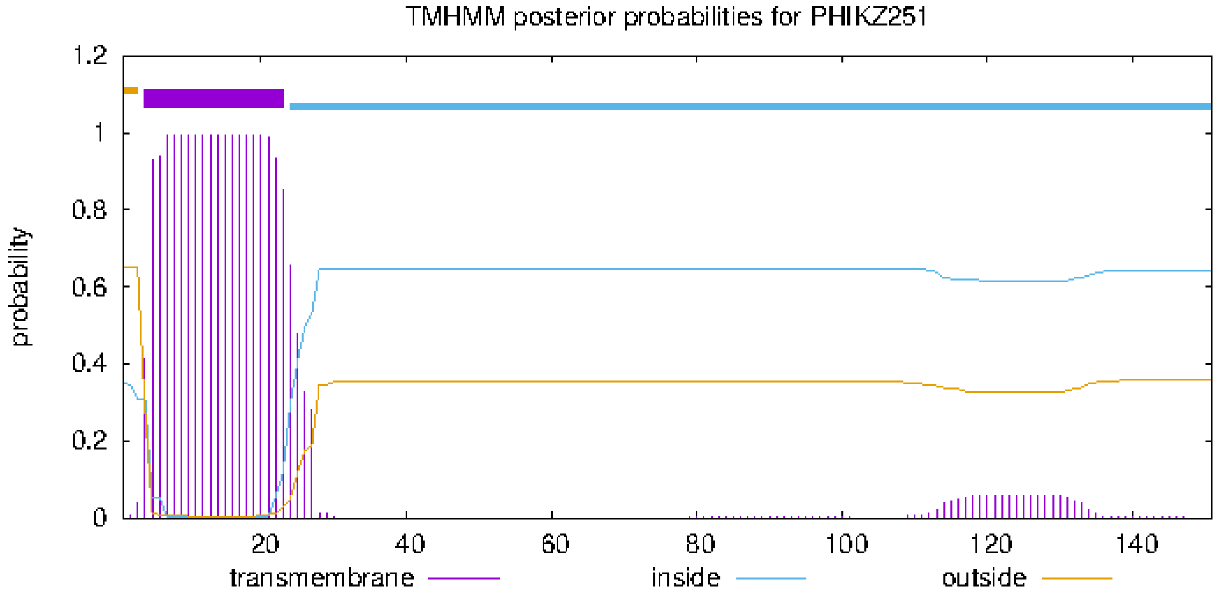


### plot in postscript, script for making the plot in gnuplot, data for plot

### PHIKZ253 Length: 79
### PHIKZ253 Number of predicted TMHs:  1
### PHIKZ253 Exp number of AAs in TMHs: 21.98033
### PHIKZ253 Exp number, first 60 AAs:  21.98033
# PHIKZ253 Total prob of N-in:        0.62087
### PHIKZ253 POSSIBLE N-term signal sequence
PHIKZ253     TMHMM2.0   outside      1     3
PHIKZ253     TMHMM2.0   TMhelix      4    26
PHIKZ253     TMHMM2.0   inside      27    79

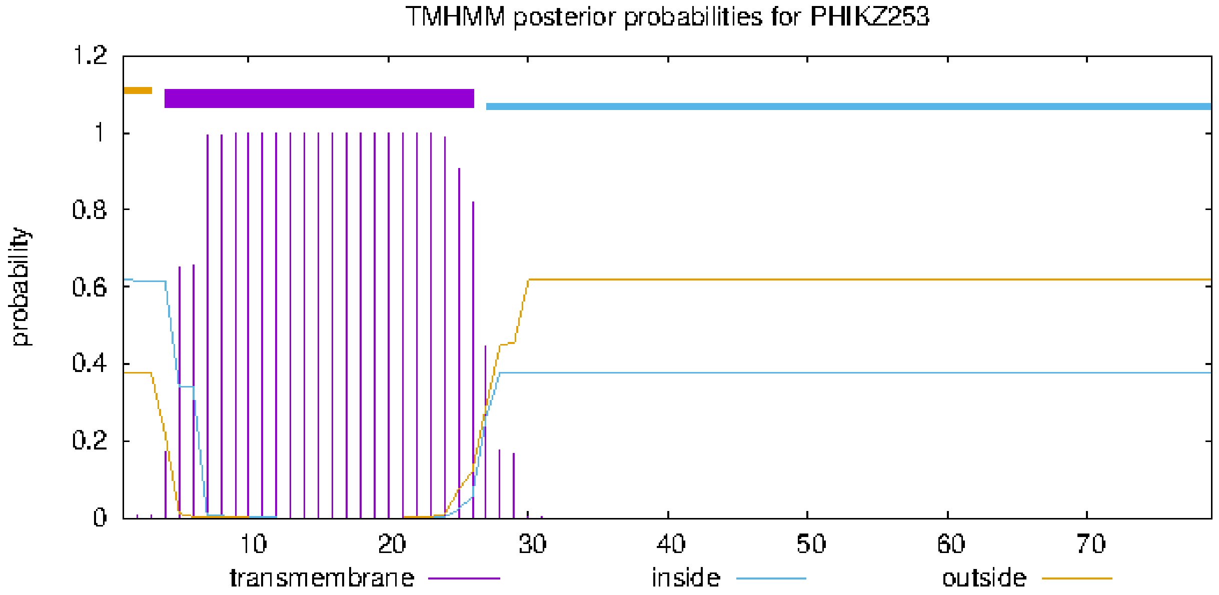


### plot in postscript, script for making the plot in gnuplot, data for plot

### PHIKZ_p74 Length: 35
### PHIKZ_p74 Number of predicted TMHs:  1
### PHIKZ_p74 Exp number of AAs in TMHs: 21.93262
### PHIKZ_p74 Exp number, first 60 AAs:  21.93262
# PHIKZ_p74 Total prob of N-in:        0.73734
### PHIKZ_p74 POSSIBLE N-term signal sequence
PHIKZ_p74   TMHMM2.0   inside      1     6
PHIKZ_p74   TMHMM2.0   TMhelix      7    29
PHIKZ_p74   TMHMM2.0   outside     30    35

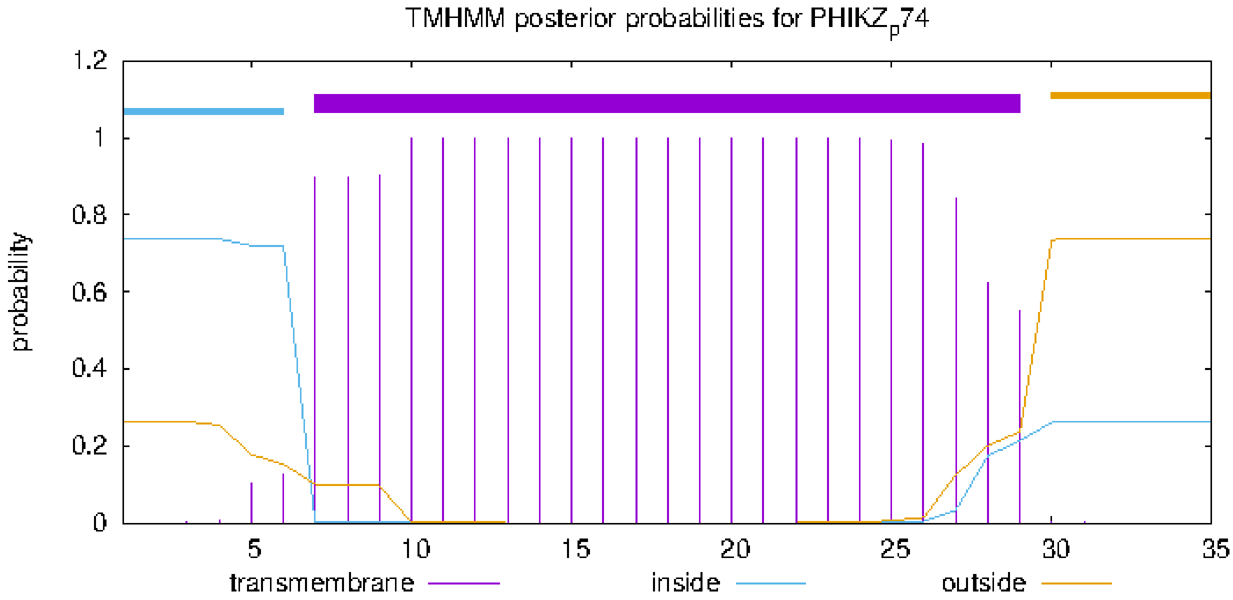


### plot in postscript, script for making the plot in gnuplot, data for plot

### PHIKZ259 Length: 144
### PHIKZ259 Number of predicted TMHs: 1
### PHIKZ259 Exp number of AAs in TMHs: 18.07914
### PHIKZ259 Exp number, first 60 AAs: 18.07869
### PHIKZ259 Total prob of N-in: 0.01147
### PHIKZ259 POSSIBLE N-term signal sequence

| | | | | |
|---|---|---|---|---|
| PHIKZ259 | TMHMM2.0 | outside | 1 | 3 |
| PHIKZ259 | TMHMM2.0 | TMhelix | 4 | 21 |
| PHIKZ259 | TMHMM2.0 | inside | 22 | 144 |

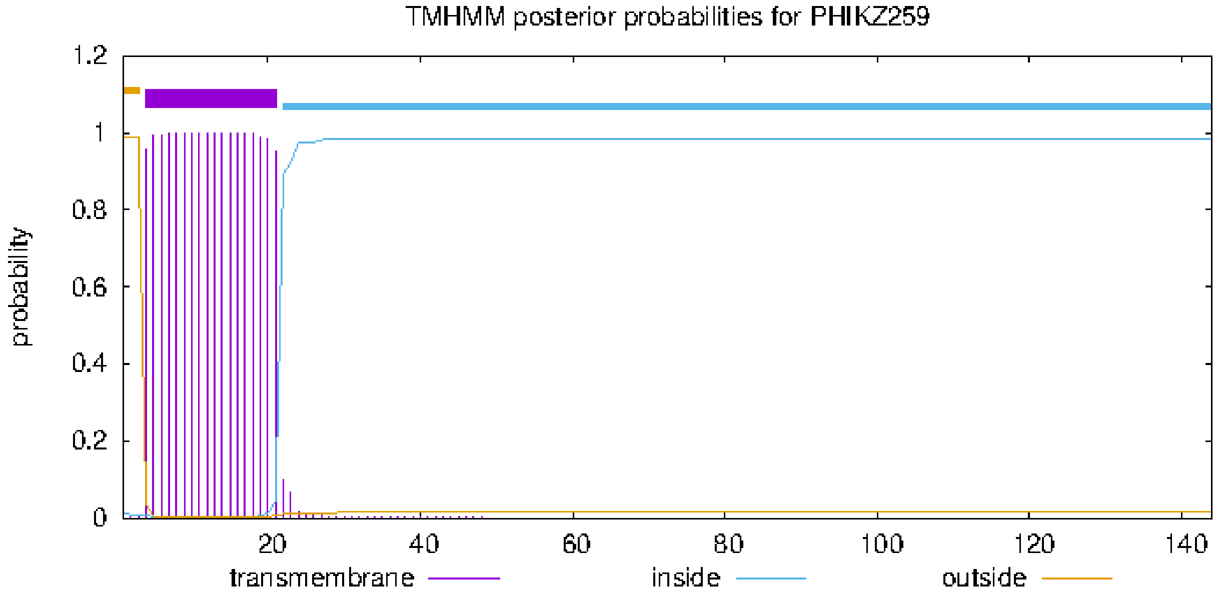


### plot in postscript, script for making the plot in gnuplot, data for plot

### PHIKZ_p77 Length: 50
### PHIKZ_p77 Number of predicted TMHs: 1
### PHIKZ_p77 Exp number of AAs in TMHs: 22.47353
### PHIKZ_p77 Exp number, first 60 AAs: 22.47353
### PHIKZ_p77 Total prob of N-in: 0.88530
### PHIKZ_p77 POSSIBLE N-term signal sequence

| | | | | |
|---|---|---|---|---|
| PHIKZ_p77 | TMHMM2.0 | inside | 1 | 6 |
| PHIKZ_p77 | TMHMM2.0 | TMhelix | 7 | 29 |
| PHIKZ_p77 | TMHMM2.0 | outside | 30 | 50 |

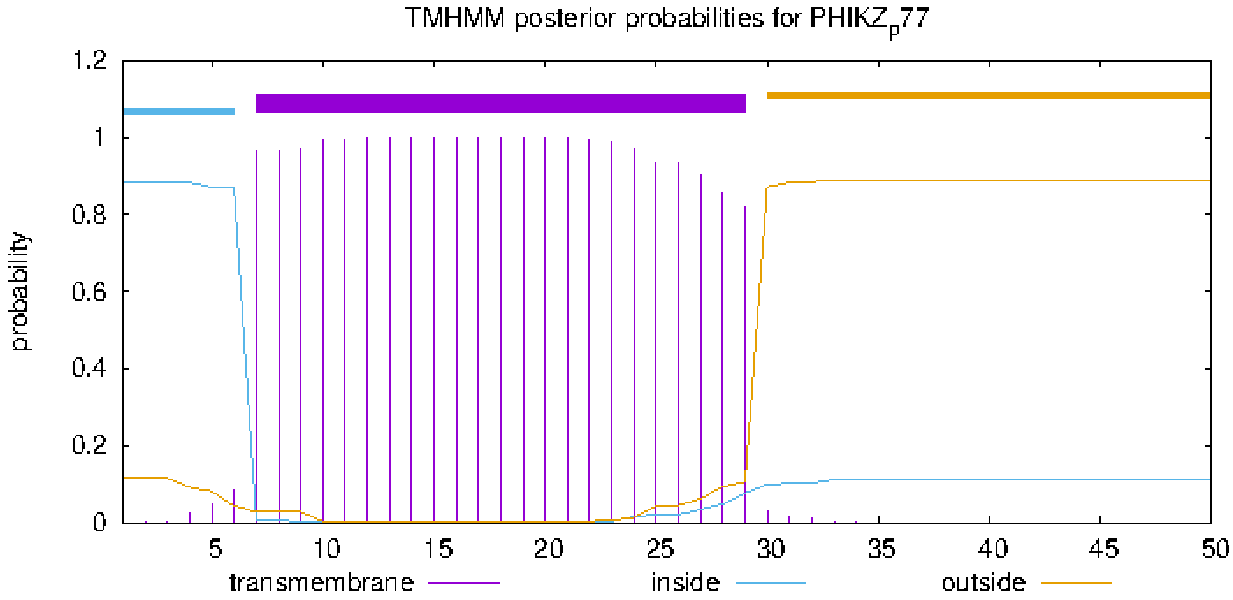


### plot in postscript, script for making the plot in gnuplot, data for plot

### PHIKZ286 Length: 508
### PHIKZ286 Number of predicted TMHs:  1
### PHIKZ286 Exp number of AAs in TMHs: 20.98447
### PHIKZ286 Exp number, first 60 AAs:  20.29564
# PHIKZ286 Total prob of N-in:        0.96763
### PHIKZ286 POSSIBLE N-term signal sequence

| | | | | |
|---|---|---|---|---|
| PHIKZ286 | TMHMM2.0 | inside | 1 | 19 |
| PHIKZ286 | TMHMM2.0 | TMhelix | 20 | 39 |
| PHIKZ286 | TMHMM2.0 | outside | 40 | 508 |

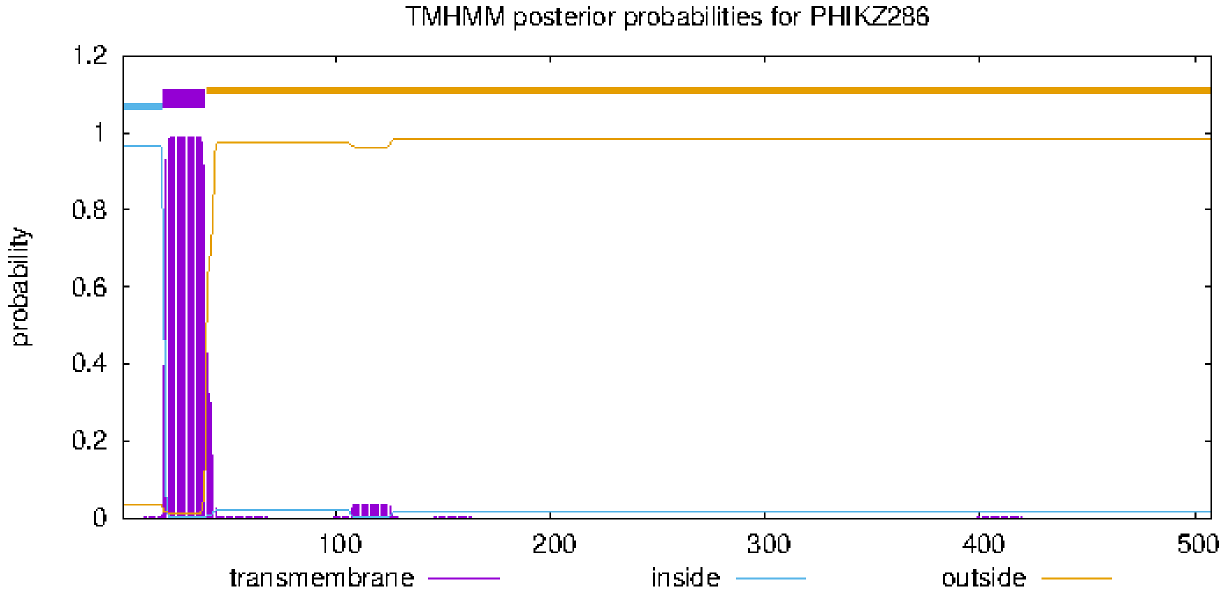


### plot in postscript, script for making the plot in gnuplot, data for plot

### PHIKZ_p82 Length: 49
### PHIKZ_p82 Number of predicted TMHs:  1
### PHIKZ_p82 Exp number of AAs in TMHs: 19.66596
### PHIKZ_p82 Exp number, first 60 AAs:  19.66596
# PHIKZ_p82 Total prob of N-in:        0.97116
### PHIKZ_p82 POSSIBLE N-term signal sequence

| | | | | |
|---|---|---|---|---|
| PHIKZ_p82 | TMHMM2.0 | inside | 1 | 26 |
| PHIKZ_p82 | TMHMM2.0 | TMhelix | 27 | 45 |
| PHIKZ_p82 | TMHMM2.0 | outside | 46 | 49 |

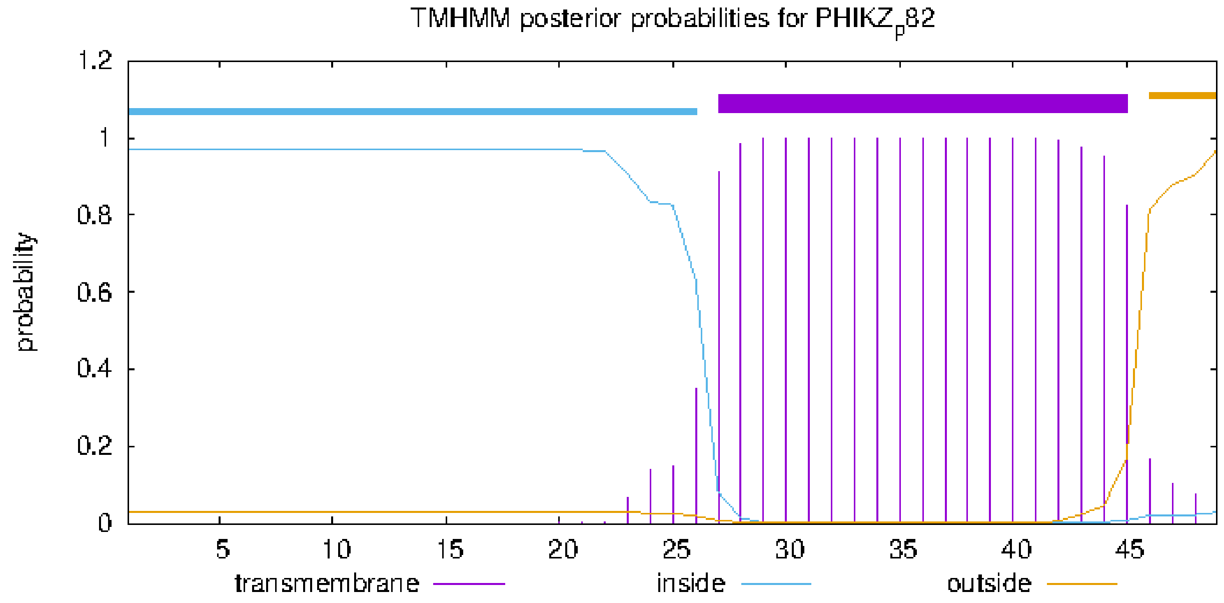


### plot in postscript, script for making the plot in gnuplot, data for plot

### PHIKZ_p91 Length: 45
### PHIKZ_p91 Number of predicted TMHs:  1
### PHIKZ_p91 Exp number of AAs in TMHs: 22.43412
### PHIKZ_p91 Exp number, first 60 AAs:  22.43412
# PHIKZ_p91 Total prob of N-in:        0.53896
### PHIKZ_p91 POSSIBLE N-term signal sequence
PHIKZ_p91   TMHMM2.0   inside      1     6
PHIKZ_p91   TMHMM2.0   TMhelix     7    29
PHIKZ_p91   TMHMM2.0   outside    30    45

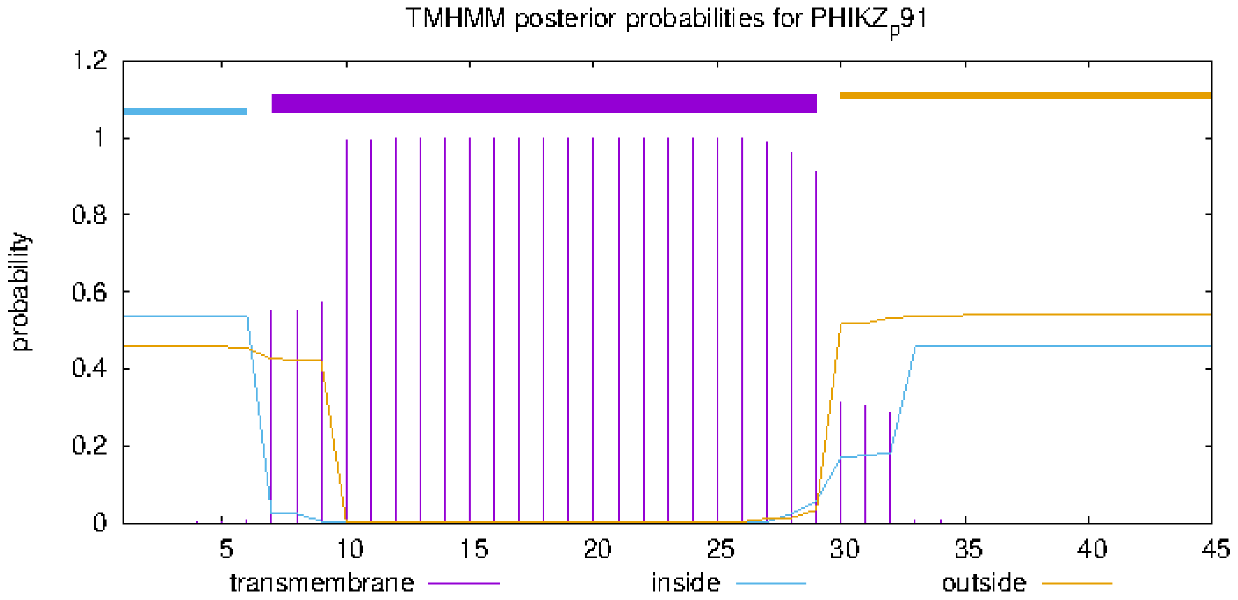


### plot in postscript, script for making the plot in gnuplot, data for plot
